# Network and State Specificity in Connectivity-Based Predictions of Individual Behavior

**DOI:** 10.1101/2023.05.11.540387

**Authors:** Nevena Kraljević, Robert Langner, Vincent Küppers, Federico Raimondo, Kaustubh R. Patil, Simon B. Eickhoff, Veronika I. Müller

## Abstract

Predicting individual behavior from brain functional connectivity (FC) patterns can contribute to our understanding of human brain functioning. This may apply in particular if predictions are based on features derived from circumscribed, *a priori* defined functional networks, which improves interpretability. Furthermore, some evidence suggests that task-based FC data may yield more successful predictions of behavior than resting-state FC data. Here, we comprehensively examined to what extent the correspondence of functional network priors and task states with behavioral target domains influences the predictability of individual performance in cognitive, social, and affective tasks. To this end, we used data from the Human Connectome Project for large-scale out-of-sample predictions of individual abilities in working memory (WM), theory-of-mind cognition (SOCIAL), and emotion processing (EMO) from FC of corresponding and non-corresponding states (WM/SOCIAL/EMO/resting-state) and networks (WM/SOCIAL/EMO/whole-brain connectome). Using root mean squared error and coefficient of determination to evaluate model fit revealed that predictive performance was rather poor overall. Predictions from whole-brain FC were slightly better than those from FC in task-specific networks, and a slight benefit of predictions based on FC from task versus resting state was observed for performance in the WM domain. Beyond that, we did not find any significant effects of a correspondence of network, task state, and performance domains. Together, these results suggest that multivariate FC patterns during both task and resting states contain rather little information on individual performance levels, calling for a reconsideration of how the brain mediates individual differences in mental abilities.

**Highlights:**

- Better prediction of behavior from task vs. resting-state FC only in a cognitive domain
- Little evidence for specificity of state, network, or task similarity
- Predicting complex behavior based on FC remains a significant challenge
- We extend research on brain-based behavior prediction beyond the cognitive domain

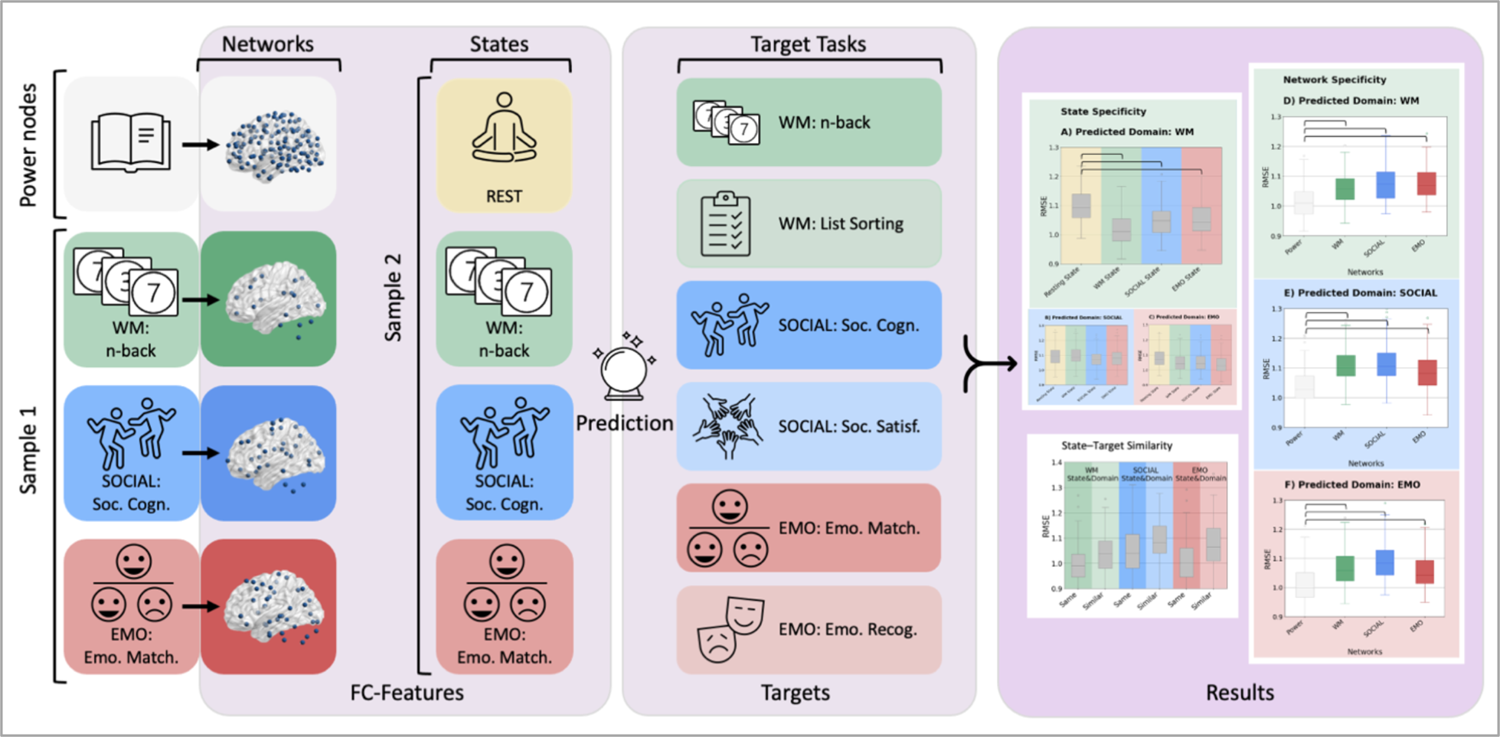

## Introduction

No two individuals are alike in perception, affect, thought and behavior, but also brain structure and function. A major goal of neuroscience is uncovering the relationships between these dimensions by investigating individual differences. An approach that has recently become popular is predicting individual behavior, affective characteristics or cognitive abilities from brain data (Gao et al., 2019; Greene et al., 2018; Kong et al., 2019; Larabi et al., 2021; Nostro et al., 2018; Ooi et al., 2022; Rosenberg et al., 2020; Sasse et al., 2022; Shen et al., 2017). Such predictive modeling is thought to yield important insights about generalizable brain–behavior relationships and is considered a crucial step towards personalized medicine (Mueller et al., 2013; D. Wang et al., 2015).

A number of studies in this area have shown that, for example, regional gray-matter volume and structural connectivity significantly predict age (Cole et al., 2017; Franke et al., 2012; More et al., 2023), reading comprehension (Cui et al., 2018), inhibitory control (N. He et al., 2020) or fear of pain (X. Wang et al., 2019). Similarly, functional neuroimaging data has also been reported to predict different behaviors or traits, ranging from personality (Dubois, Galdi, Han, et al., 2018; Nostro et al., 2018) or life satisfaction (Itahashi et al., 2021) to cognitive abilities such as creative thinking (Zhuang et al., 2021), cognitive flexibility (Chén et al., 2019), or working memory (WM) capacity (Stark et al., 2021).

While a variety of different brain characteristics have been employed as features to predict behavior, one of the most widely used measures (Yeung et al., 2022) is resting-state (in the following also “rest” or “REST”) functional connectivity (FC) obtained from functional magnetic resonance imaging (fMRI). Some recent studies, however, suggest that behavior prediction may benefit from the use of *task-based* FC, as compared to resting-state data (Avery et al., 2020; Greene et al., 2018; Jiang et al., 2020; Rosenberg et al., 2016, 2016; Stark et al., 2021). For example, Sripada and colleagues found that the correlation between predicted and observed scores of a general cognitive ability factor improved when using FC from the 2-back WM-task state (r = 0.50), as compared to using resting-state FC (r = 0.26; (Sripada et al., 2019, 2020). A similar pattern has been reported for the prediction of different measures of attention (Yoo et al., 2018) as well as for the prediction of intelligence based on FC from tasks taxing executive functioning (L. He et al., 2021) or attention (Rosenberg et al., 2016).

Importantly, all the studies mentioned above showed an improvement in prediction performance for task-fMRI data derived from the same domain as the predicted measure (i.e. prediction of stop-signal task performance based on FC derived from stop-signal task-fMRI data). However, there is not only resting versus task states but rather different task states depending on which task is performed during fMRI data acquisition. That is, every task performed in the scanner can be thought of as eliciting a specific state. Interestingly, it has been shown that in predicting intelligence, using almost any other task state (i.e. fMRI acquired during a WM task as well as an emotion task) or task–rest combinations outperforms using resting-state FC only (Gao et al., 2019; Greene et al., 2018, 2020; Sripada et al., 2020).

Based on the concept of convergent and discriminant validity (Campbell & Fiske, 1959; Schumann et al., 2022), it would be expected, however, that connectivity patterns observed during the same or a similar task, hence coming from the same domain as the predicted target behavior (i.e., representing the same state; convergent validity), lead to better prediction performance than do patterns observed during a task state from an unrelated domain (discriminant validity). In line with this idea, recent studies reported better accuracies for predicting general cognitive ability (Sripada et al., 2020) and fluid intelligence (Gao et al., 2019) from FC during task states involving executive functions (“same-domain”), as compared to prediction from unrelated task or resting states (“other-domain”). This improvement was particularly pronounced when FC data of the cognitively demanding WM task was used, as compared to task states from other domains (although the authors did not test for the statistical significance of the observed numerical differences between prediction accuracies). These examples suggest the possibility of state specificity when predicting behavior from corresponding FC patterns.

While most studies predicted task performance from states of the same domain (i.e. prediction of intelligence from FC of a WM task state), others predicted task performance from FC observed during the exact same task. Avery et al. (2020), for example, predicted individual performance accuracy in an n-back WM task based on FC derived from fMRI data obtained while the n-back task was performed, which showed an increase in accuracy when using this task’s fMRI data, as compared to rest data (Avery et al., 2020). Building on this, Stark and colleagues investigated the difference in prediction accuracy between predictions of performance in different working and episodic memory tasks from FC obtained while performing an n-back WM task. Importantly, the highest prediction accuracy (r = 0.36) was achieved when n-back task performance was predicted using FC during the very same task (i.e. n-back performance measured in the MR scanner), followed by the prediction of performance in a different WM task (list sorting; r = 0.24), followed by predictions of episodic memory performance scores (r = 0.05-0.11) (Stark et al., 2021). These results suggest a specificity benefit of the state used for calculating FC, which goes beyond the prediction of ability in a given broadly defined cognitive domain (i.e. WM) and narrows it down to specific tasks (i.e. n-back versus list sorting). That is, beyond the effect of *state* specificity (i.e., domain congruence benefits for prediction accuracy), a *state–target* similarity effect (i.e., task congruence benefits for prediction accuracy) should manifest in even better prediction accuracies for the performance in tasks during which FC data were acquired, as compared to the performance in other tasks of the same domain.

The literature to date is inconclusive regarding the effects of state specificity and state–target similarity on FC-based predictions of mental abilities and psychological traits. In particular, the majority of studies investigating such prediction models have focused on cognitive targets such as intelligence or attention (Yeung et al., 2022). Thus, clear evidence for state specificity and state– target similarity is still lacking, especially in domains like emotion processing and social cognition.

The studies above mainly used whole-brain FC for behavior prediction. Sometimes, *post-hoc* examination of the most predictive features from the whole-brain feature space are used for better interpretability (e.g.: (Chén et al., 2019; Dubois, Galdi, Paul, et al., 2018; Itahashi et al., 2021; Jiang et al., 2020; Pläschke et al., 2020). However, such *post-hoc* analyses come with their own limitations, as feature weights are context-dependent, their reliability is rather low, and the results can be highly specific to the given dataset (Tian & Zalesky, 2021). Besides predicting behavior from whole-brain FC, several studies reported on predictions using particular functional networks as priors (J. Chen et al., 2021; Heckner et al., 2023; Nostro et al., 2018). That is, prediction models in these investigations exclusively rely on FC between regions that show activation during a given task. It is argued that this aides and constrains the functional interpretability of any observed associations (e.g. most predictive features), since such models are based on brain regions for which the association between brain and mental function has already been established independently.

Therefore, network-based prediction has the advantage of better interpretability of the results due to the *a priori* knowledge about the mental function a given network subserves (Nostro et al., 2018; Pläschke et al., 2017). Similar to state specificity, FC within networks associated with functions that are more closely related to the target behavior (e.g., predicting WM performance from WM network features) should also be more informative than networks that are associated with very different functions (e.g., predicting WM performance from pain network features). Few studies have investigated this network specificity, with some suggesting some network specificity with regard to personality (Nostro et al., 2018), but others showing a lack of specificity (Heckner et al., 2023; Pläschke et al., 2020).

The current project, therefore, aimed to investigate the influence of brain state (same-vs other-domain), similarity of target behavior to the features within one domain (same vs. similar task from same domain), and functional network priors (same-vs other-domain network) on the predictability of individual behavior. This included the specific question of whether FC from same-domain states and in same-domain networks can predict individual behavior better than FC from other-domain states or networks. Hence, we tested the following three hypotheses: (1) State specificity: behavior should be better predicted based on FC patterns observed in the same domain, hence during the state corresponding to the behavior to be predicted, as compared to FC patterns observed in other (non-corresponding) domains. (2) State–target similarity: task performance should be better predicted based on FC patterns observed during the exact same task, as compared to another similar task from the same domain. (3) Network specificity: behavior should be better predicted based on FC patterns observed in the networks corresponding to the predicted behavior, as compared to FC patterns in other (non-corresponding) networks.

## Methods

To investigate whether there is state specificity, state–target similarity and/or network specificity in brain–behavior prediction, we used the Human Connectome Project (HCP) Young Adult dataset. We divided it into two samples: in the first sample we defined networks, and in the second sample we computed FC in predefined networks from the first sample during different task states. Using FC within each network as features, we predicted six different target variables, matching the selected states and networks. We included the following three phenotypic domains: working memory (WM), theory of mind/social cognition (SOCIAL), and emotion processing (EMO).

### Samples

Data was obtained from the Young Adult S1200 release of the publicly available database provided by the HCP (Van Essen et al., 2013), which comprised data from 1206 healthy individuals. We only included participants for whom all the data required for our analyses were available. That is, (a) all 4 resting-state fMRI scans, (b) fMRI data of the WM, SOCIAL, and EMO tasks, and (c) the performance measures (accuracy and reaction time) of these three tasks performed in the scanner, as well as (d) all the performance measures we aimed to predict for tasks that were performed outside the scanner for each domain. Hence, every subject was required to have both (in-scanner and out-of-scanner) tasks per domain (3 domains, 6 tasks in total). Of the 1206 individuals, 180 participants were excluded due to missing imaging data and 71 due to data quality issues. We further excluded subjects with accuracy below 50% in the 6 tasks of interest (n = 77). Performance accuracy was measured as the percentage of correct trials. We chose to include only subjects producing more than 50% correct trials, to ensure that only participants were included who were attentive during the task and hence present the given states we aimed to investigate. From the remaining sample of 878 subjects, two non-overlapping sub-samples were randomly generated: one for independently delineating task-based networks via general linear modelling (GLM; Sample 1) and one for brain–behavior prediction within those (and other) networks (Sample 2). Thus, the first sample can be thought of as the sample for “network extraction” and the second sample for “feature extraction and prediction”. We carefully accounted for the family structure and resulting dependencies in this dataset by ensuring that (i) sample 1 contained only one individual per family, (ii) there was no kinship between the two samples, and (iii) a leave-n-family-out cross-validation (CV) scheme for prediction analyses within sample 2 was used (Poldrack et al., 2020). For an overview of the sample selection, see **Error! Reference source not found.**.

Sample 1, used for delineating task networks, consisted of 250 unrelated subjects (138 females; age mean = 28.6 years, standard deviation (SD) = 3.8, range = 22–36 years). Sample 2, used for prediction, consisted of all the remaining individuals with no siblings in the first sample. Further, we removed individuals that scored higher/lower than four standard deviations from the mean in any of the six target scores of interest, leaving us with 467 participants (252 females, mean age = 28.8 years, standard deviation (SD) = 3.7, range = 22–36 years) for sample 2. Note that this sample contains siblings, which was accounted for in the prediction pipeline through family sub-sampling in the CV (Poldrack et al., 2020). From sample 2 we further randomly selected a hold-out sample (47 subjects), which was not used in any of the CVs. Therefore, sample 2 consisted of 420 individuals that were used for CV and final training, while n = 47 participants were held back for subsequently testing generalizability. The analyses of the HCP data were approved by the ethics committee of the Medical Faculty at the Heinrich Heine University Düsseldorf.

### Network delineation

Two different approaches were employed for delineating task-specific networks: (i) networks reflecting brain activation in a large sample of participants during the tasks of interest using the task fMRI data of sample 1, and (ii) activation likelihood estimation (ALE) meta-analyses across previously published neuroimaging results of the same tasks. For brevity, we here only report the methods and results of the first approach to network delineation. Further details on the results of the second, meta-analytic, approach can be found in the supplementary material.

### Delineation of task-networks in Sample 1

Ultimately, our network extraction approach aimed to delineate networks that were as closely as possible related to the states we aimed to predict in the second sample. For this, we included strictly only the task of interest for network delineation in both approaches – the single study and the meta-analyses. To cover a variety of domains, we chose three very different tasks performed in the scanner: n-back for the WM domain, emotion recognition / face processing for the EMO domain, and social cognition / theory of mind for the SOCIAL domain. For details on the tasks, see (Barch et al., 2013). Briefly, an n-back task was used for WM, presenting a sequence of different stimuli with the instruction to either decide whether the current stimulus is the same as the one used 2 trials ago (2-back) or to recognize a specific target (0-back). EMO was a face matching task in which angry or fearful faces had to be matched (EMO condition), in contrast to matching shapes (neutral condition). In the SOCIAL task, animated moving shapes were shown in either interacting or random manner and had to be labelled subsequently as interacting or randomly moving.

For the delineation of the task-specific networks revealing nodes that are activated in our task of interest, we used the minimally preprocessed volumetric task fMRI data of participants of sample 1. The preprocessing included artifact removal, motion correction, and registration to the MNI standard volume space. More details regarding the preprocessing pipeline can be found in Glasser et al., 2013. The minimally preprocessed data was input for the general linear modelling (GLM), performed using FSL (Version 5.0.9) (Jenkinson et al., 2012; Smith et al., 2004; Woolrich et al., 2009). For the subject-level GLM we modified the scripts provided by the HCP ((Barch et al., 2013); https://github.com/Washington-University/HCPpipelines), which are based on the FSL FEAT module (Woolrich et al., 2001, 2004), for use of volumetric data.

The subject-level GLM included for either run (two different phase encoding directions) temporal high-pass filtering (200 s cutoff), spatial smoothing (8 mm FWHM Gaussian kernel), and the GLM fitting. The respective stimuli in each task were modeled as blocked predictors, temporal derivatives of each predictor, 6 movement parameters and their derivatives as regressors of no interest. For each task, linear contrasts between conditions were computed: 2-back > 0-back for WM; interaction > random for SOCIAL; and faces > shapes for EMO, respectively. Data across both phase encoding directions were then combined with a fixed-effects GLM analysis.

For the group-level GLM, we modified an FSL workflow developed by (Esteban et al., 2019), which estimated the group effects using FSL FLAME (FMRIB’s Local Analysis of Mixed Effects) by performing a one-sample t-test across subjects. Group-level activation maps were thresholded at cluster-level p < 0.05 (FWE-corrected for multiple comparisons) with a cluster-forming threshold of p < 0.001. From the resulting clusters, we extracted only the peak coordinates with a minimum distance of 15 mm. This resulted in three networks: WM-NW, SOCIAL-NW, and EMO-NW. For an overview of the workflow, please see Figure 2. For comparison with the task-specific networks, we used FC between the Power nodes (Power et al., 2011) as a functionally defined, spatially distributed, whole-brain representation of the connectome. The Power nodes represent a combination of resting-state FC ROIs and task-based meta-analytic ROIs, yielding 264 non-overlapping independent ROIs.

**Figure 1.**
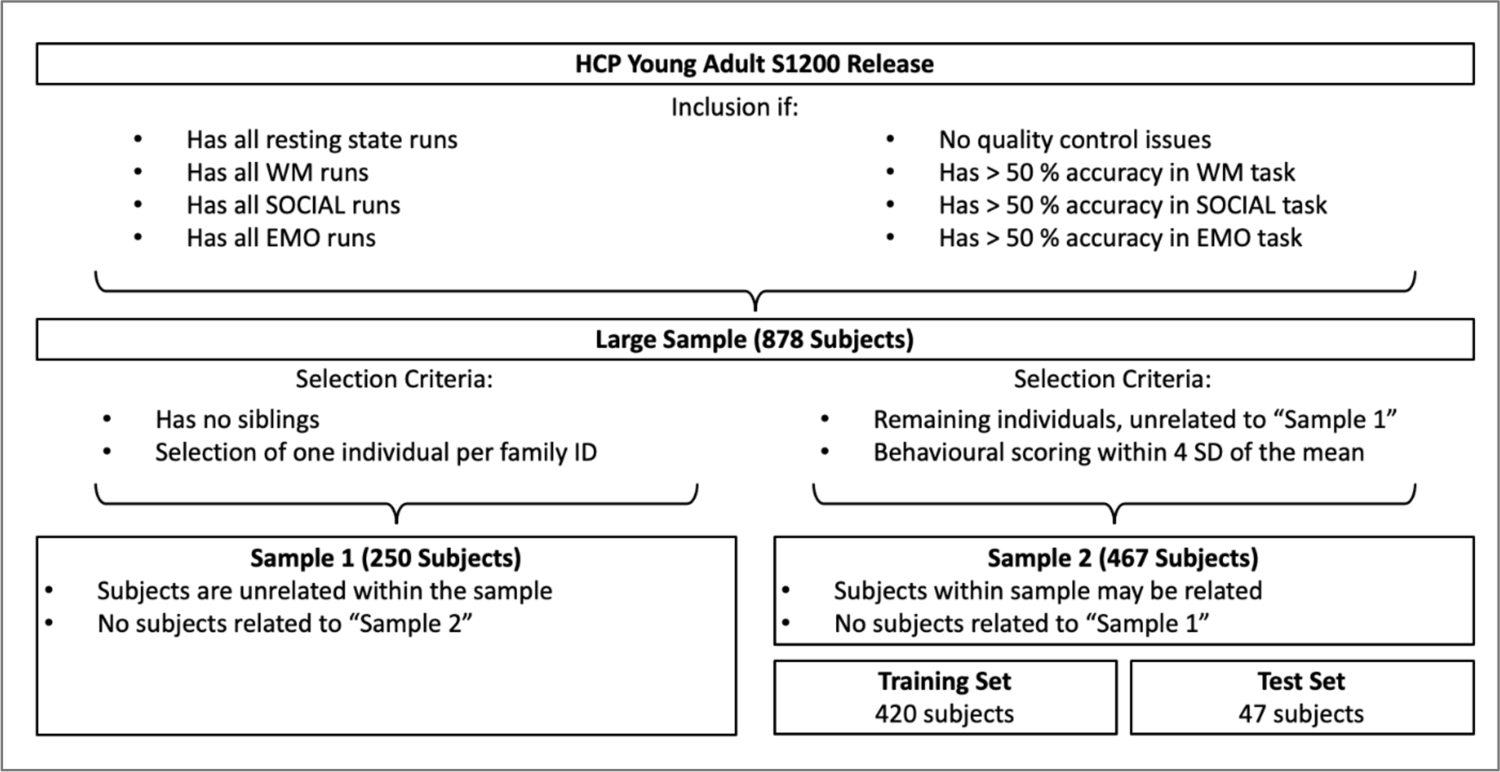
Overview of the sampling procedure.

**Figure 2.**
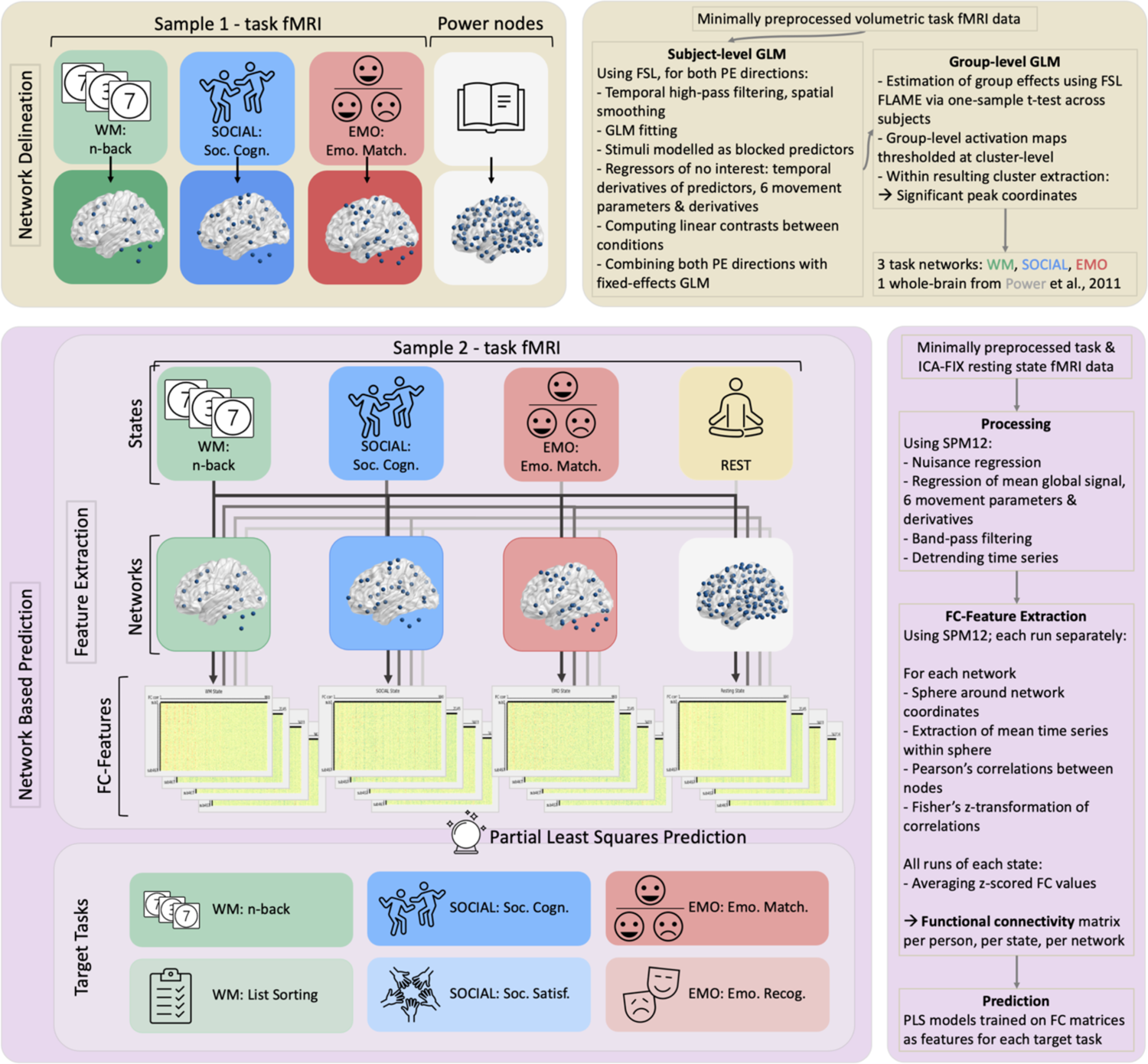
Overview of the applied methods. Yellow blocks depict the network extraction from sample 1. Violet blocks depict the network based prediction in sample 2, together with the feature extraction (functional connectivities) from the networks delineated in the first step and sample. The upper heatmap under “FC-Features” shows the FC from the different states in the WM-network. the GLM: General linear modelling, PE: Phase encoding, FC: Functional connectivity, Soc. Cog.: Social cognition task (in-scanner task), Soc. Satisf.: Social Satisfaction Questionnaire (out-of-scanner score), Emo. Match.: Emotional Face Matching task (in-scanner task), Emo. Recog.: Emotional Face Recognition task (out-of-scanner score).

Finally, we compared the resulting task-networks from both sample 1 and the meta-analyses, to ensure they covered and showed overlap with frequently observed regions in previously reported large-scale analyses (WM: Daamen et al., 2015; Fuentes-Claramonte et al., 2019; Kennedy et al., 2017; Rottschy et al., 2012; Snoek et al., 2021; https://identifiers.org/neurovault.collection:7103; SOCIAL: T. Chen et al., 2023; Hennion et al., 2016; Mossad et al., 2022; Patil et al., 2017; EMO: Chaudhary et al., 2023; Herrmann et al., 2020; Nord et al., 2017; Snoek et al., 2021; https://identifiers.org/neurovault.collection:7103). The resulting WM-NW had 49 nodes, the SOCIAL-NW had 71 nodes, and EMO-NW had 84 nodes. For an overview of the networks, please see Figure 2. For a complete view of the networks and further details on the exact coordinates, please refer to supplemental Figures S1-7 and the supplemental Tables S2-4.

### Prediction in sample 2

#### Targets: Behavioral Measures

To assess state specificity and state–target similarity, we selected behavioral performance during different tasks: First, we used performance collected in the scanner for our three domains of interest (WM, SOCIAL, EMO → “same task”/ in-scanner task). Second, we selected scores of tasks/questionnaires that measured behavior not exactly in the same state but still in the same behavioral domains (“similar task” / out-of-scanner task). The two levels of tasks (“same” and “similar”) from the same domain enable us to advance insights beyond state specificity, into state– target similarity.

For “same task” scores (in-scanner task), reaction time and accuracy of task performance were used. These two scores were combined by calculating the Inverse Efficiency Score (IES; (Townsend & Ashby, 1983)), which is defined as the mean response time across correct trials of the condition of interest divided by its accuracy. This was employed to address the issue of ceiling effects in the accuracy scores. Hence, for WM (subsequently called “n-back”), IES was calculated using mean response time and accuracy of the 2-back blocks. For EMO, we used response time and accuracy in the face-block of the emotional face matching task (subsequently called “matching” or “EMO matching”). For SOCIAL, since the accuracy in both interaction and random trials involved theory-of-mind cognition (Castelli et al., 2000), we averaged response time and accuracy of both interaction and random trials before creating the IES (subsequently called “Social Cognition”).

For “similar task” scores (out-of-scanner task or questionnaire scores) in the WM domain, we selected the unadjusted list sorting score from the NIH Toolbox List Sorting Working Memory Test (subsequently called “List Sorting”). For SOCIAL, we computed a social satisfaction compound score (Babakhanyan et al., 2018) across five different scales (friendship, loneliness, emotional support, instrumental support, and perceived rejection) of the self-report Emotion Battery of the NIH Toolbox (Salsman et al., 2013) (subsequently called “Social Satisfaction” or “Satisfaction”). For EMO, we computed the IES using reaction time and accuracy of the Penn Emotion Recognition Test (Gur et al., 2001, 2010) (subsequently called “Emotion Recognition” or “Recognition”). See supplementary Table 1 for an overview of all targets included.

#### Features: Functional connectivity

Resting-state fMRI and the three sets of task fMRI data (WM, SOCIAL, EMO) from sample 2 were used for calculating FC within each network of interest (WM, SOCIAL, EMO and Power). The network extraction is explained in detail in the section “Delineation of task-networks in Sample 1”; for an illustration of the methods applied, please see Figure 2. For all four states we used all runs available (4 runs for REST and 2 runs each for the tasks) and their full duration per run. MRI protocols of HCP were previously described in detail (Glasser et al., 2013; Van Essen et al., 2013). For the task-fMRI data, we used the minimally preprocessed version provided by the HCP, which includes removal of spatial distortions, volume realignment, registration to the anatomical image, bias field reduction, normalization to the global mean, and masking the data with the final brain mask (Glasser et al., 2013). The approach to treat task fMRI comparable to resting state fMRI data has been suggested by (Greene et al., 2020). For the resting-state fMRI data, we used the ICA-FIX denoised data provided by the HCP, which uses the minimally preprocessed fMRI data (processed in the same way as task fMRI data) as input and denoises it through classification of ICA components. This classifier identifies “good” and “bad” components and automatically removes artifactual or “bad” components. For further details see (Griffanti et al., 2014; Salimi-Khorshidi et al., 2014; Smith et al., 2013).

Additional processing as well as the functional connectivity analysis for both resting-state and task-fMRI data were performed using SPM12 (www.fil.ion.ucl.ac.uk/spm/software/spm12/) and MATLAB 2020a (The Mathworks, Natick, MA). Nuisance regression was done to control for mean white-matter and cerebrospinal-fluid signals, mean global signal, within-scanner motion using the 6 movement parameters and their derivatives stored in the Movement_Regressors_dt.txt file provided by the HCP. Further, we applied band-pass filtering [0.01-0.1 Hz] and detrended the time series. We opted for the band-pass filter, as this has been shown to be successful in filtering out movement and physiological artefacts, without leading to information loss (Ciric et al., 2017; Satterthwaite et al., 2013). Using the network coordinates obtained from sample 1 (depicted in Figure 2 and in the supplemental material in Figures S1-7 and Tables S2-4), for each network, we modeled a 5-mm sphere around each node’s coordinate. The sphere size was the same for all coordinates to ensure the same number of voxels within each node. However, as we extracted multiple peak coordinates from larger clusters of task-activation in sample 1, larger activated regions are represented by multiple spheres. From each sphere, we extracted the mean time series. We then calculated the Pearson’s correlations between all pairs of nodes of each respective network, before applying Fisher’s Z-transformation. These steps were done for each run separately (4 runs for REST and 2 runs each for the tasks) and for each state (REST, WM, SOCIAL, and EMO). The z-scored FC values were finally averaged across all runs of each state. This was done for all 4 networks as well as for each of the four states. Connectivity matrices for the WM network can be seen in Figure 2, all other networks can be found in the supplemental material (Figures S8-14).

To ensure that any effects were not due to the different lengths of the tasks performed in the scanner, we trimmed all time series to the length of the shortest scan duration (EMO: 2:16 mins) for a control analysis. These results are reported in the supplemental material (Figure S16).

#### Network-based prediction of individual behavior

We predicted the task performance/characteristics for each domain from resting- and task-state FC (4 states: REST, WM, EMO, SOCIAL; to investigate state specificity) and task of interest (same and similar tasks in WM, EMO, SOCIAL; to further investigate state–target similarity) of the delineated networks (4 networks: whole-brain Power nodes and 3 task networks (WM, SOCIAL, EMO from sample 1; to investigate network specificity).

For the main analysis we used Partial Least Squares (PLS) as the prediction model. PLS is a form of supervised learning which uses linear regression fitting, but it can handle violation of the assumption of no multicollinearity by reducing the dimensionality of correlated variables. However, to confirm our results and to cover models that have been used in the past for behavioral prediction, we additionally performed analyses using other algorithms: Kernel Ridge Regression, which, like PLS, is a linear parametric model. As well as Support Vector Regression (with both linear and non-linear RBF kernel) and Random Forest as nonparametric models, where both can capture non-linear relationships. Lastly, we used PLS and kernel ridge also with connectivity-based prediction modelling (CBPM; Finn et al., 2015; Shen et al., 2017) as a popular feature reduction. CBPM correlates the features to the target variable, retaining only the features showing a significant relation to the target for the model to learn. Note that all models were trained using the same set of FC features and target variables. All results from the additional analyses are presented in the supplemental material (Figures S17-22).

Separate prediction analyses were conducted for each combination of network, state, and behavioral score, resulting in a total of 4 states x 4 networks x 6 targets = 96 predictions. The FC pattern of the respective network and state constituted the given feature space, and the respective behavioral scores were the targets. All algorithms were used as implemented in JuLearn (Hamdan et al., 2023), which is a toolbox based on scikit-learn (Pedregosa et al., 2011). It includes hyperparameter tuning, nested-CV, and feature reduction methods making sure that data leakage is avoided. For PLS, we tuned the hyperparameters in an inner 5-fold CV, with the number of latent components increasing in steps from 1 to 10. As having a sibling in the training set could lead to a better prediction of the related participant’s score in the validation set, we applied a 100 x leave-30%-families-out CV scheme on 420 subjects from sample 2 to account for the family structure of the sample (i.e. individuals from the same family were not split into training or validation sample but kept in either one of them). This is done to counter potential nonindependence induced by the family structure in the HCP dataset (Poldrack et al., 2020). We deconfounded the features by regressing out age and sex as well as normalizing them by z-transformation. For comparability of prediction performance between the different behavioral scores, we additionally normalized the targets. To avoid data leakage, confound regression and normalization were done within the CV. That is, the confound regression models and parameters for z-transformation were computed in the CV on the training set only (70% of the families) and then applied to the test set (30% of the families) (Poldrack et al., 2020). Prediction performance was evaluated by the root mean squared error (RMSE) as well as the coefficient of determination (COD), as a measure of goodness of model fit, averaged across all CV runs. Additionally, the mean Pearson correlation across all CV runs between predicted and observed scores was calculated. After hyperparameter tuning and CV, we finally applied the model, that has been trained on all the data provided and with the hyperparameter tuning performed on it, to the randomly drawn hold-out sample (47 subjects from sample 2) to evaluate the model’s generalizability.

The RMSE was used for testing for significant differences of prediction performance between states, networks and, tasks using machine-learning (ML)-adjusted t-tests (Nadeau & Bengio, 1999). These modified t-tests are evaluated and adjusted for comparing machine-learning algorithms (Bouckaert & Frank, 2004) to account for violating the independence assumption in a paired Student’s t-test. This is done by correcting the variance estimate through considering the training and sample size. In our case, due to the leave-30%-families-out CV scheme, the number of data points changed in each fold. Therefore, we used the mean training sample size across the 100 folds for the adjustment.

Within each phenotypic domain, we first tested effects of state and network by averaging prediction performance of the respective other factors (i.e., averaging across networks and task when testing for state effects, and across state and task when testing for network effects). As state–target similarity is an extension of state specificity, we here only averaged across networks for same and similar tasks, respectively. Significant effects (Bonferroni corrected for multiple comparisons) were then further assessed by comparing the respective individual prediction scores between each other.

## Results

We mainly report on the outcomes of the prediction analyses using PLS. Predictions using different algorithms and approaches showed highly similar patterns and their details can be found in the supplementary material. Furthermore, regarding outcome measures, we here focus on the RMSE from the CV as well as the coefficient of determination (COD) as a measure of goodness of fit. In the supplementary material, additional results of Pearson’s r of the predicted and observed score from the CV can be found.

Averaged across all CV-folds per prediction, the COD and RMSE (Figure 3) revealed that the models show a poor fit and prediction accuracies are rather low. Note, that COD values below zero indicate that prediction of individual scores were worse than predicting the mean of the target. The mean COD showed a positive mean value only for 2 out of 96 predictions, while all others showed a mean COD of zero or a negative value. Other models (e.g. kernel ridge regression or SVR) yielded some more COD values above zero, but no model achieved a mean COD higher than 0.07. Similarly, the RMSE was quite high for all predictions. Because these scores indicated a generally poor fit to the data, we refrained from applying the best model to the hold-out sample. The correlations (for details, see the supplementary material) between predicted and observed values ranged from −0.11 to 0.32 with a mean prediction accuracy of 0.08 (standard deviation: 0.09) and only one mean correlation from the 96 predictions reaching a medium effect size.

**Figure 3.**
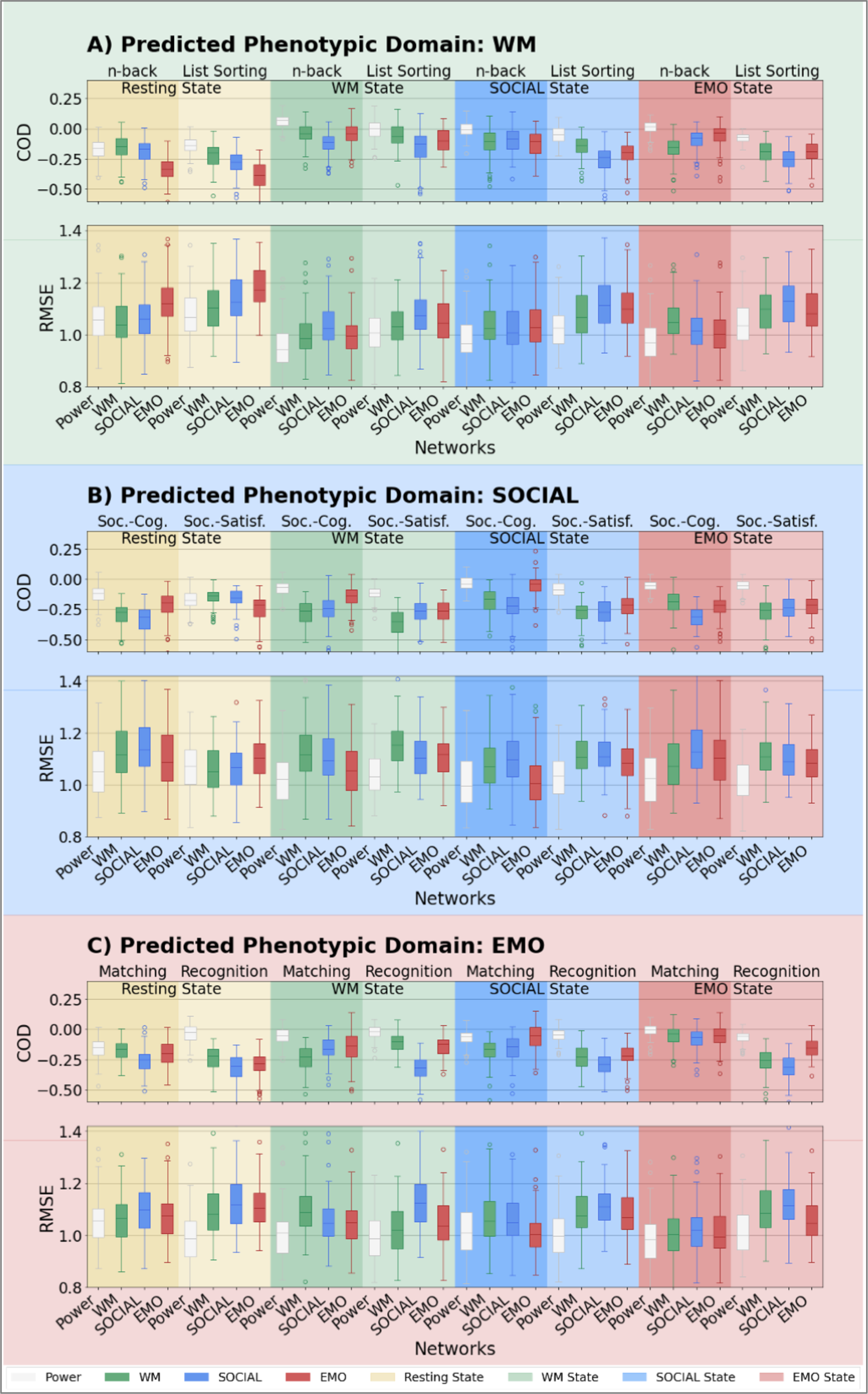
Prediction performance: Boxplots of the distribution of COD and RMSE from the 100 x CV for each phenotypic domain (A – WM, B – Social, C – EMO domain), state and network. Bars represent the model fit / COD (all negative values are set to 0) and RMSE of prediction of a specific score (WM, SOCIAL, EMO; performed in (darker background) or outside (lighter background) the scanner) based on functional connectivity within a given network (POWER, WM, SOCIAL, EMO) in a given state (REST, WM, SOCIAL, EMO). Green: WM, blue: SOCIAL, red: EMOTION, yellow: resting state, white: Power nodes. Darker background: target is the task performed in the scanner; lighter background: target is the task performed outside the scanner. Soc. Cog.: Social cognition task (in-scanner task), Soc. Satisf.: Social Satisfaction Questionnaire (out-of-scanner score), Matching: Emotional Face Matching task (in-scanner task), Recognition: Emotional Face Recognition task (out-of-scanner score).

### State specificity in network-based prediction

To answer the question if the correspondence between state and target (e.g. WM score predicted using FC during WM state) improves predictions, and whether there even is state specificity (e.g. WM scores predicted significantly better from FC during WM state than from FC during other states: REST, SOCIAL, or EMO), we examined the differences in prediction accuracy between states.

No significant differences were found for the SOCIAL and EMO domain (Figure 4 B-C). For the WM domain, the ML-adjusted t-test (Bonferroni corrected for multiple comparisons) showed a significant benefit for all task states compared to the resting state (see Figure 4-A; see Table 1 for mean RMSE and significant t-test statistics). However, non-WM domain states (i.e., SOCIAL and EMO) only showed a significant difference to the WM state at an uncorrected threshold (not shown in Table 1). This difference was also significant when using other algorithms and feature selection approaches (PLS with CBPM, Random Forest, SVR with the RBF kernel). At an uncorrected threshold, the difference was also significant for all other models (kernel ridge regression, SVR with linear kernel, and CBPM with ridge regression), as well as when using only the trimmed time-series.

**Figure 4.**
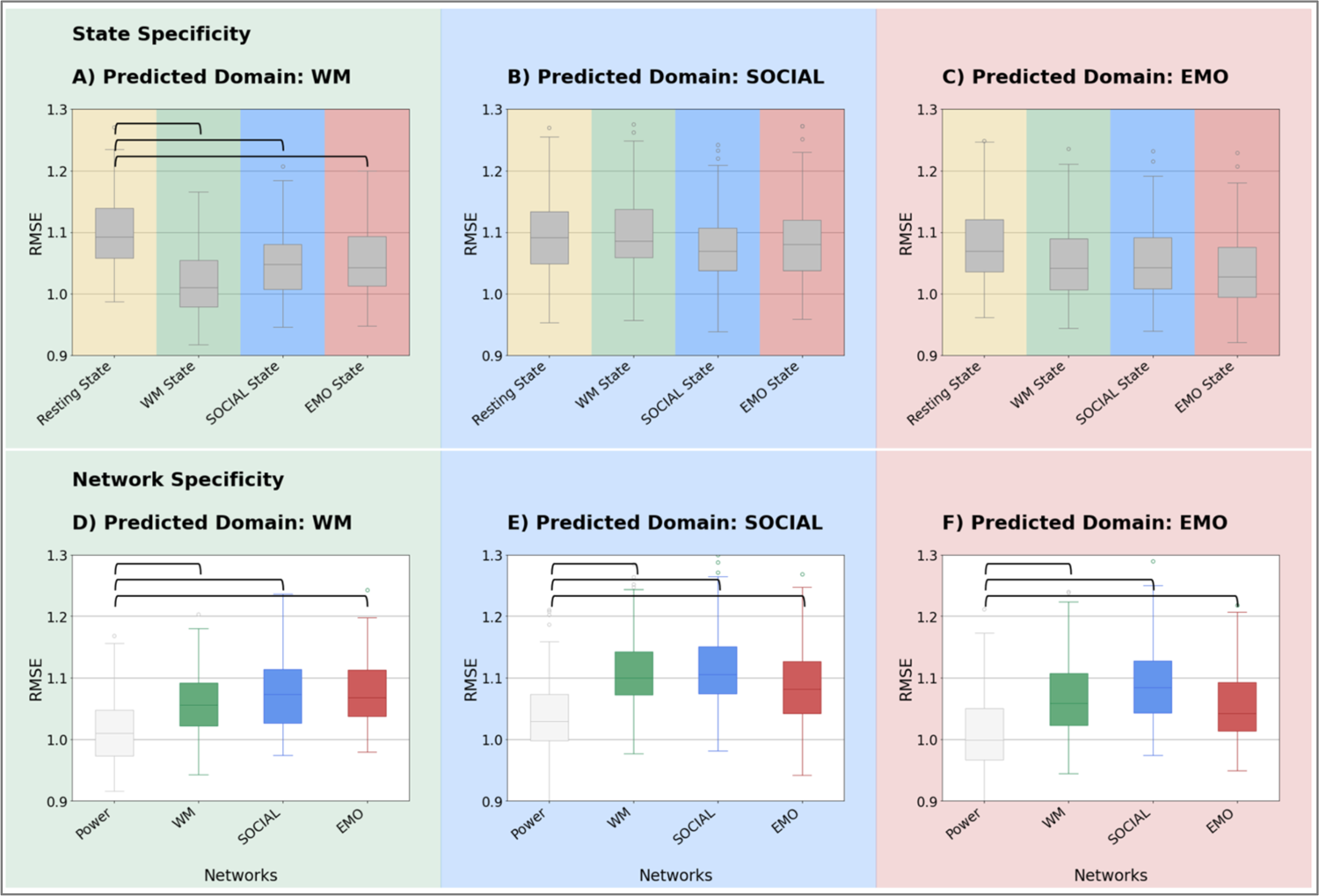
State and network specificity: State (A-C) and network (D-E) specificity for each phenotypic domain (A and D – prediction of WM scores, B and E – prediction of SOCIAL scores, C and F – prediction of EMO scores). For state specificity (A-C) RMSE of all networks (POWER, WM, SOCIAL, EMO) and the two tasks of the respective domain in a given state (REST, WM, SOCIAL, EMO) are averaged. For network specificity all states (REST, WM, SOCIAL, EMO) and the two task of the respective domain are averaged in a given network (Power nodes, WM, SOCIAL, EMO). Green: WM, blue: SOCIAL, red: EMOTION, yellow: resting state. Horizontal bars indicate significance p_corr_ < 0.05.

**Table 1.** Comparison of prediction accuracies between states.

| Domain of predicted performance | State A | RMSE mean (SD) | State B | RMSE mean (SD) | *t* | *p*-value |
| --- | --- | --- | --- | --- | --- | --- |
| **Significant main effect of state** |  |  |  |  |  |  |
| **WM** | REST | 1.10 (0.06) | WM | 1.02 (0.05) | 5.10 | < 0.001 |
|  | REST | 1.10 (0.06) | SOCIAL | 1.05 (0.06) | 3.49 | 0.013 |
|  | REST | 1.10 (0.06) | EMOTION | 1.05 (0.06) | 3.68 | 0.007 |
| **Significant *post-hoc* tests for specific state—task—network combinations** |  |  |  |  |  |  |
| **WM** | REST (n-back, Power nodes) | 1.06 (0.09) | WM (n-back, Power nodes) | 0.95 (0.09) | 4.83 | < 0.001 |
|  | REST (n-back, EMO network) | 1.13 (0.10) | WM (n-back, EMO network) | 1.00 (0.10) | 3.51 | 0.016 |
|  | REST (List Sorting, EMO network) | 1.19 (0.09) | WM (List Sorting, EMO network) | 1.05 (0.09) | 4.10 | 0.002 |
|  | REST (n-back task, Power nodes) | 1.06 (0.09) | SOCIAL (n-back task, Power nodes) | 0.98 (0.09) | 3.21 | 0.043 |
|  | REST (n-back task, Power nodes) | 1.06 (0.09) | EMOTION (n-back task, Power nodes) | 0.97 (0.09) | 3.72 | 0.008 |
|  | REST (n-back, EMO network) | 1.13 (0.10) | EMOTION (n-back, EMO network) | 1.00 (0.10) | 3.40 | 0.023 |
|  | REST (List Sorting, EMO network) | 1.19 (0.09) | EMOTION (List Sorting, EMO network) | 1.10 (0.09) | 3.32 | 0.030 |

*Note.* Machine-learning-adjusted t-test to assess state specificity using the averaged 100 RMSE values obtained from 100-fold cross-validation within the state listed in column “State A” versus the state listed in column “State B”. *p*-values are Bonferroni corrected for multiple comparisons. *Post-hoc* t-tests between individual predictions of the task in the network (both noted in brackets) and the state listed in column “State A” versus the state listed in column “State B”.

To asses which effect was driving the significant differences we compared prediction performance between states using *post-hoc* ML-adjusted t-tests, while keeping network and task constant. That is, we only compared predictions between same-domain networks and tasks (e.g. comparing the prediction performance of “same” WM task score based on FC within Power nodes in *resting state* to the prediction performance of “same” WM task score based on FC within Power nodes in *WM state*). Comparing prediction performance between rest and different states for each network and WM score revealed that the difference between REST and WM state was driven by the difference in prediction performance of the n-back task using the Power nodes and the EMO network (for mean RMSE and significant t-test statistics, see Table 1). The EMO network additionally showed a difference between WM and REST state when predicting list sorting. The difference between REST and EMO state was also driven by Power nodes and the EMO network when predicting n-back performance, and by the EMO network when predicting list sorting. The significant difference in REST versus SOCIAL was driven by the Power nodes in predicting n-back performance.

Overall, state had a significant influence on predicting WM scores, with better predictions when using task compared to resting state. This state-specific improvement was, however, not uniformly observed and mainly driven by predictions based on FC within the Power nodes and the EMO network.

### State–target similarity in network-based prediction

In a next step, we examined differences in predictability between the “same” and “similar” tasks in a given domain. We were interested in whether the behavior would be predicted better in the state where the predicted behavior was measured (“same task”), and whether the FC-based predictivity could translate to a related task (“similar task”). An example would be the comparison between the predictability of n-back task performance (WM task performed during scanning) and the list sorting task (WM task performed outside of the scanner) based on FC patterns observed during the WM n-back task.

For this comparison of “same task” with “similar task”, we found a slight (numerical) benefit in the performance of the “same task”. However, a direct statistical comparison of RMSE values using ML-adjusted t-tests did not show any significant effects after Bonferroni correction for multiple comparisons (see Figure 5).

**Figure 5.**
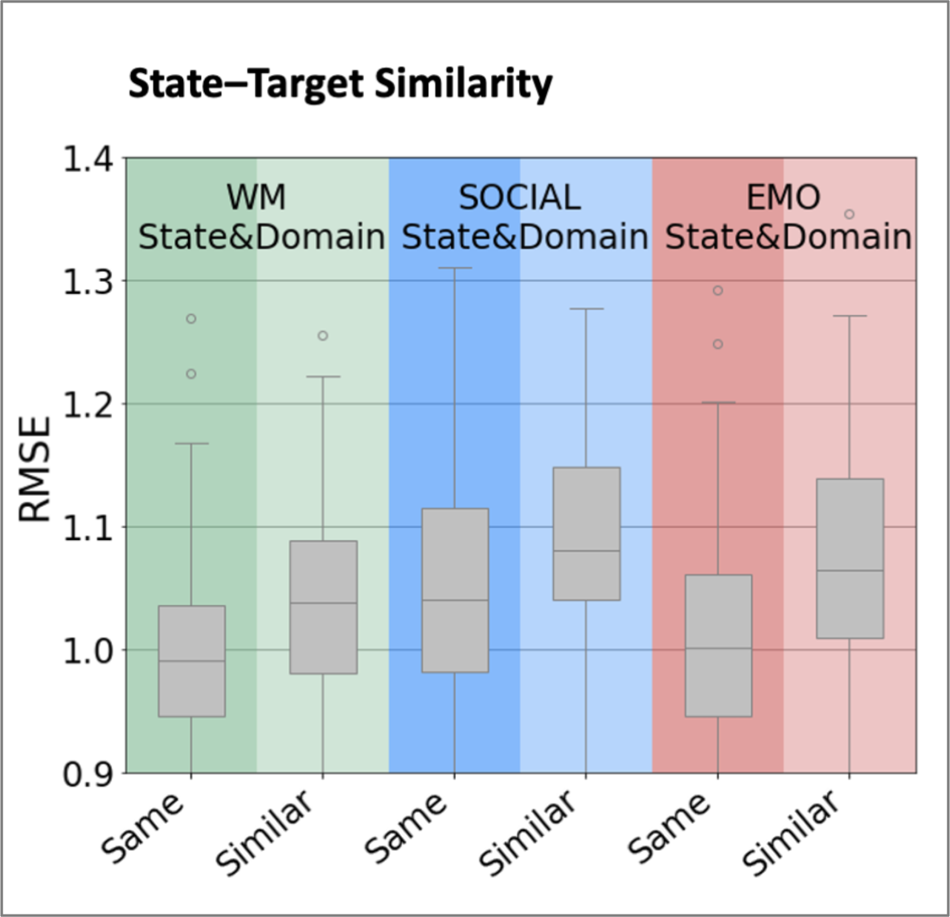
State–target similarity: Boxplots of the distribution of RMSE RMSE from the 100 x CV averaged across all networks (Power nodes, WM, SOCIAL, EMO network) within a given state (WM, SOCIAL and EMO) and task (SAME or SIMILAR) Green: WM, blue: SOCIAL, red: EMOTION, grey: averaged across networks. Darker background: target is the “same” task performed in the scanner, lighter background: target is the “similar” task performed outside the scanner.

### Network specificity in network-based prediction

Finally, we set out to answer the question if predicting task performance does benefit from being based on FC within a network known to be engaged in performing that same task (e.g. n-back task performance predicted from FC within the WM-network), as compared to other task-related networks (e.g. n-back task performance predicted from FC within a SOCIAL task-based network or the whole-brain connectome). ML-adjusted t-tests showed, for all three domains, a benefit for the whole-brain Power nodes over the task-specific networks (see Figure 4 D-F). In the WM domain, FC between the Power nodes predicted WM-related targets better than did FC within the n-back WM-specific network, the SOCIAL-specific network, or the EMO-specific network (see Figure 4-D and Table 2 for mean RMSE and t-test statistics). In the SOCIAL domain, the Power nodes predicted SOCIAL-related targets better than did the WM-specific, SOCIAL-specific, or EMO-specific networks (see Figure 4-E). Finally, in the EMO domain, the Power nodes again predicted EMO-related targets better than all three domain-specific networks including the EMO-specific network derived from an emotional face-matching task (see Figure 4-F).

**Table 2.**
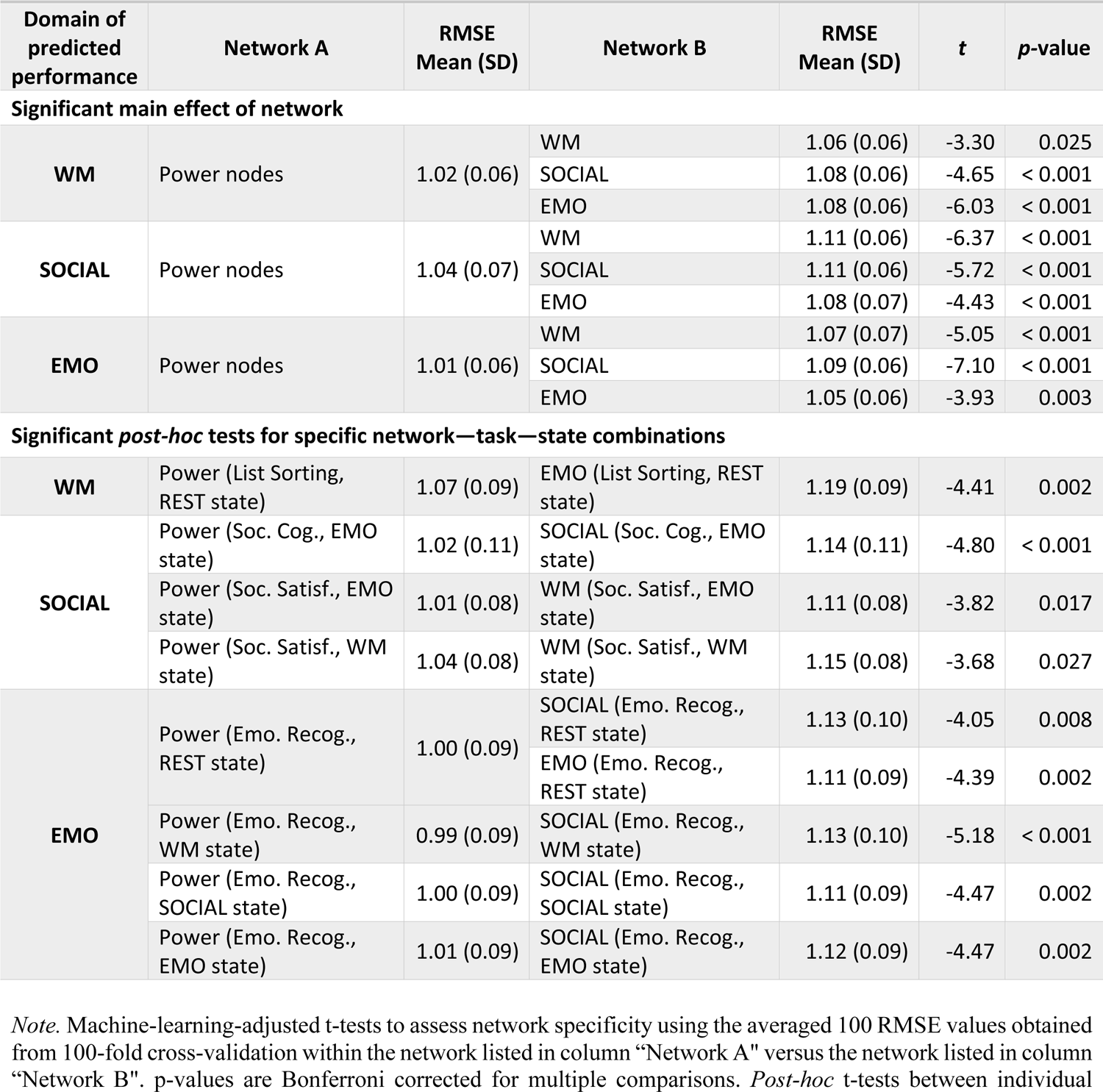
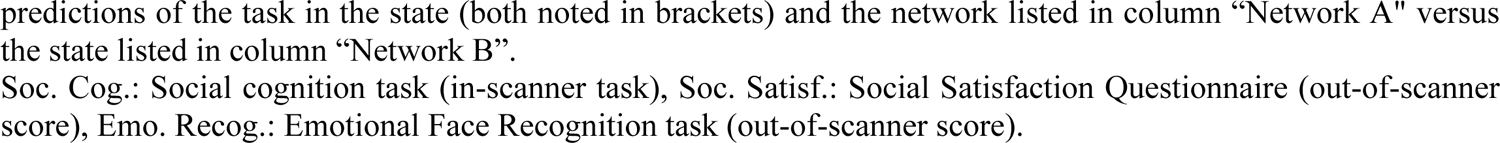
Comparison of prediction accuracies between networks.

| Domain of predicted performance | Network A | RMSE Mean (SD) | Network B | RMSE Mean (SD) | *t* | *p*-value |
| --- | --- | --- | --- | --- | --- | --- |
| Significant main effect of network |  |  |  |  |  |  |
| WM | Power nodes | 1.02 (0.06) | WM | 1.06 (0.06) | -3.30 | 0.025 |
|  |  |  | SOCIAL | 1.08 (0.06) | -4.65 | < 0.001 |
|  |  |  | EMO | 1.08 (0.06) | -6.03 | < 0.001 |
| SOCIAL | Power nodes | 1.04 (0.07) | WM | 1.11 (0.06) | -6.37 | < 0.001 |
|  |  |  | SOCIAL | 1.11 (0.06) | -5.72 | < 0.001 |
|  |  |  | EMO | 1.08 (0.07) | -4.43 | < 0.001 |
| EMO | Power nodes | 1.01 (0.06) | WM | 1.07 (0.07) | -5.05 | < 0.001 |
|  |  |  | SOCIAL | 1.09 (0.06) | -7.10 | < 0.001 |
|  |  |  | EMO | 1.05 (0.06) | -3.93 | 0.003 |
| Significant *post-hoc* tests for specific network—task—state combinations |  |  |  |  |  |  |
| WM | Power (List Sorting, REST state) | 1.07 (0.09) | EMO (List Sorting, REST state) | 1.19 (0.09) | -4.41 | 0.002 |
| SOCIAL | Power (Soc. Cog., EMO state) | 1.02 (0.11) | SOCIAL (Soc. Cog., EMO state) | 1.14 (0.11) | -4.80 | < 0.001 |
|  | Power (Soc. Satisf., EMO state) | 1.01 (0.08) | WM (Soc. Satisf., EMO state) | 1.11 (0.08) | -3.82 | 0.017 |
|  | Power (Soc. Satisf., WM state) | 1.04 (0.08) | WM (Soc. Satisf., WM state) | 1.15 (0.08) | -3.68 | 0.027 |
| EMO | Power (Emo. Recog., REST state) | 1.00 (0.09) | SOCIAL (Emo. Recog., REST state) | 1.13 (0.10) | -4.05 | 0.008 |
|  |  |  | EMO (Emo. Recog., REST state) | 1.11 (0.09) | -4.39 | 0.002 |
|  | Power (Emo. Recog., WM state) | 0.99 (0.09) | SOCIAL (Emo. Recog., WM state) | 1.13 (0.10) | -5.18 | < 0.001 |
|  | Power (Emo. Recog., SOCIAL state) | 1.00 (0.09) | SOCIAL (Emo. Recog., SOCIAL state) | 1.11 (0.09) | -4.47 | 0.002 |
|  | Power (Emo. Recog., EMO state) | 1.01 (0.09) | SOCIAL (Emo. Recog., EMO state) | 1.12 (0.09) | -4.47 | 0.002 |

*Note.* Machine-learning-adjusted t-tests to assess network specificity using the averaged 100 RMSE values obtained from 100-fold cross-validation within the network listed in column “Network A” versus the network listed in column “Network B”. *p*-values are Bonferroni corrected for multiple comparisons. *Post-hoc* t-tests between individual

**Table 3.**
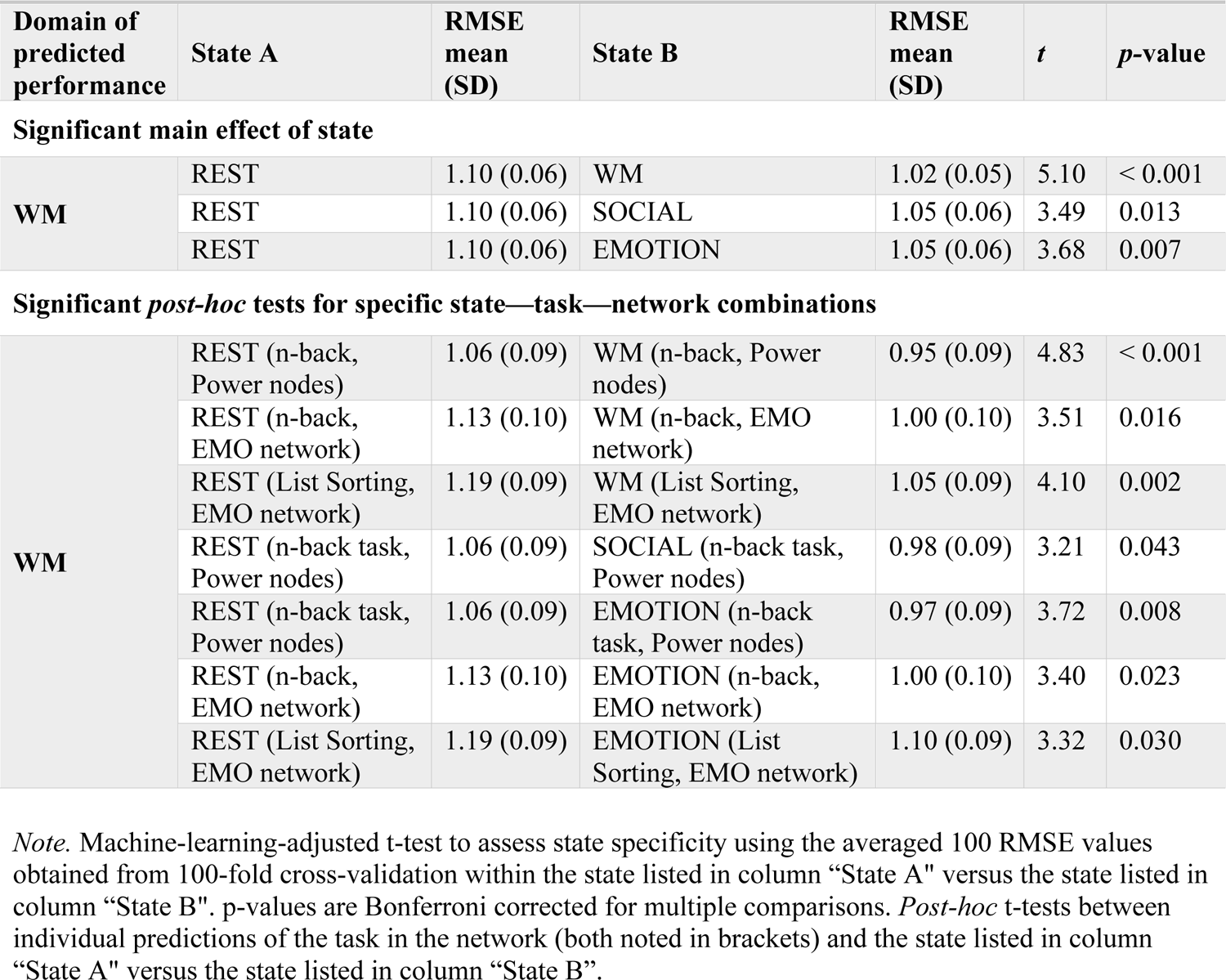
Comparison of prediction accuracies between states.

| Domain of predicted performance | State A | RMSE mean (SD) | State B | RMSE mean (SD) | *t* | *p*-value |
| --- | --- | --- | --- | --- | --- | --- |
| Significant main effect of state |  |  |  |  |  |  |
| WM | REST | 1.10 (0.06) | WM | 1.02 (0.05) | 5.10 | < 0.001 |
|  | REST | 1.10 (0.06) | SOCIAL | 1.05 (0.06) | 3.49 | 0.013 |
|  | REST | 1.10 (0.06) | EMOTION | 1.05 (0.06) | 3.68 | 0.007 |
| Significant *post-hoc* tests for specific state—task—network combinations |  |  |  |  |  |  |
| WM | REST (n-back, Power nodes) | 1.06 (0.09) | WM (n-back, Power nodes) | 0.95 (0.09) | 4.83 | < 0.001 |
|  | REST (n-back, EMO network) | 1.13 (0.10) | WM (n-back, EMO network) | 1.00 (0.10) | 3.51 | 0.016 |
|  | REST (List Sorting, EMO network) | 1.19 (0.09) | WM (List Sorting, EMO network) | 1.05 (0.09) | 4.10 | 0.002 |
|  | REST (n-back task, Power nodes) | 1.06 (0.09) | SOCIAL (n-back task, Power nodes) | 0.98 (0.09) | 3.21 | 0.043 |
|  | REST (n-back task, Power nodes) | 1.06 (0.09) | EMOTION (n-back task, Power nodes) | 0.97 (0.09) | 3.72 | 0.008 |
|  | REST (n-back, EMO network) | 1.13 (0.10) | EMOTION (n-back, EMO network) | 1.00 (0.10) | 3.40 | 0.023 |
|  | REST (List Sorting, EMO network) | 1.19 (0.09) | EMOTION (List Sorting, EMO network) | 1.10 (0.09) | 3.32 | 0.030 |

*Note.* Machine-learning-adjusted t-test to assess state specificity using the averaged 100 RMSE values obtained from 100-fold cross-validation within the state listed in column “State A” versus the state listed in column “State B”. *p*-values are Bonferroni corrected for multiple comparisons. *Post-hoc* t-tests between individual predictions of the task in the network (both noted in brackets) and the state listed in column “State A” versus the state listed in column “State B”.

**Table 4.**
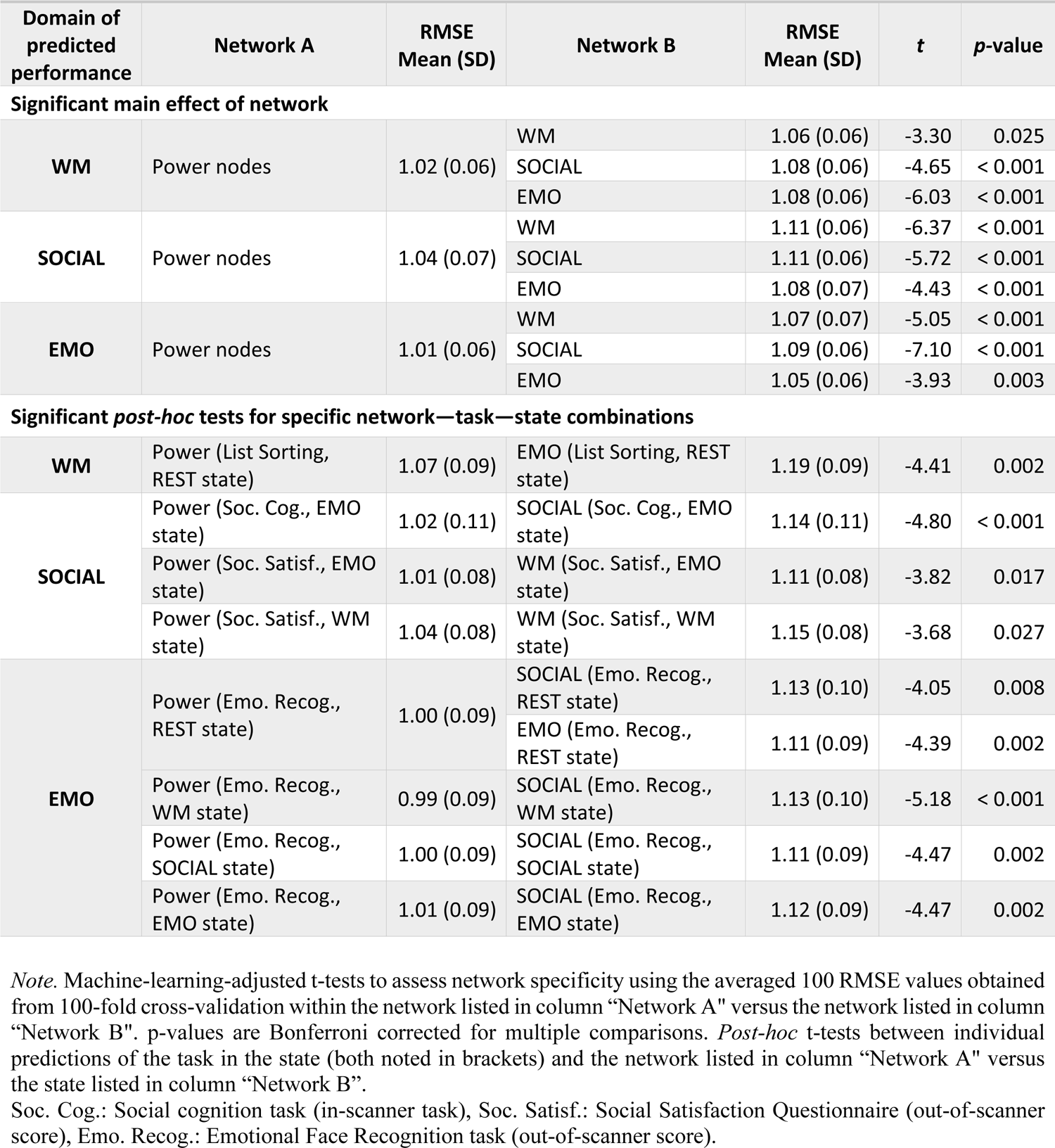
Comparison of prediction accuracies between networks.

*Post-hoc* tests (see Table 2) indicated that the difference between the Power and WM networks was driven by the EMO and WM state when predicting social satisfaction, while there was no specific state or task driving the effects for the EMO and WM domain. The priority of Power over the SOCIAL-specific network was in particular evident when predicting social cognition and social satisfaction in the EMO state and when predicting emotion recognition in the REST, SOCIAL, and EMO states. No specific state or task was driving the effect for the WM domain. The effect of Power versus the EMO-specific network was driven by the predictions of list sorting and emotion recognition in the REST state. No specific state or task was driving the effect in the SOCIAL domain.

This superiority of the Power nodes over functional network definitions in all three domains was not present in other ML algorithms. However, we still saw a similar trend when trimming the time series to the shortest task (EMO: 2:16 mins), as the Power nodes performed better in all domains and networks, except for the EMO domain where the EMO network did not perform significantly worse.

## Discussion

Using state-of-the-art fMRI preprocessing and ML approaches, this study investigated brain– behavior relationships. Specifically, how brain features from specific states and networks, or the task similarity within the behavioral domain, affects this relationship. Based on previous studies, we hypothesized that brain features obtained from networks and/or states that are corresponding to the target task are more informative about individual behavior than those obtained from other (non-corresponding) states or networks. Additionally, we expected that behavior in the task performed during fMRI data acquisition will be predicted better than similar tasks of the same phenotypic domain. Contrary to expectations, we found no significant differences in predictability (when correcting for multiple comparisons) that would indicate specific benefits of state, task, or network correspondence. Rather, our results show a general benefit of predicting WM scores using (any) task state, relative to rest, and for predicting performance in any domain from whole-brain FC (Power nodes), relative to predefined functional networks. Importantly, however, prediction accuracies were overall quite low, raising the question to what extent the observed differences (or their absence) in prediction performance between state and network conditions, and task similarity can be meaningfully interpreted.

### Is there state specificity for brain–behavior prediction?

We expected not only an improvement in predicting behavior based on FC during task states compared to FC at rest as demonstrated by (Greene et al., 2018), but especially in predicting behavior based on FC within corresponding states. However, apart from the overall low prediction performance, our results, show only weak evidence for the former, i.e. an advantage of task states compared to resting state, but only for predicting WM scores (see Figure 4-A). Previous studies have already reported that predictions of cognitive scores such as intelligence or attention improve when using task fMRI data (vs. resting state) to derive FC patterns (Avery et al., 2020; Greene et al., 2018; Jiang et al., 2020). But also when combining task and rest (Jiang et al., 2020), and specifically when using FC within a WM task state (Avery et al., 2020; Jiang et al., 2020; Sripada et al., 2020; Stark et al., 2021). Our results extend this work to different states and additionally show that this benefit of using task-fMRI data cannot be assumed for behaviors other than WM. Task-fMRI data may lead to better predictions compared to resting-state data, particularly for WM performance, possibly because task-based fMRI has a more constrained setup which potentially enhances reliability. Resting-state fMRI has been shown to be less reliable than stimulating (such as movies or task) fMRI (Noble et al., 2019). Task-based modulations of brain states may therefore contain more information about individual differences in brain functioning and behavior (Greene et al., 2018). This is consistent with a recent emphasis on shifting from solely focusing on resting-state FC (Finn, 2021; Greene et al., 2018) to accelerate progress in human neuroscience. Possibly, predictive performance can be improved by using naturalistic stimuli (Finn & Bandettini, 2021), as such settings are still more constrained than rest but less constrained than certain laboratory tasks. However, using movie data would not readily allow testing of state specificity effects, and therefore would not be a ready-made solution to the current research question.

### Is there state–target similarity?

Surprisingly, prediction accuracy was low even when FC was derived from the exact same task state in which the behavioral data was collected, with no improvement when predicting the same score, as compared to a similar score (see Figure 5). Also using the HCP dataset and WM task data, Stark et al. (2021) reported similar though slightly higher accuracies than ours for n-back (“same task”) and list-sorting (“similar task”) scores, with higher correlations for the former (but not tested for significance). We here extend these insights by also testing effects of task similarity for SOCIAL and EMO scores and by showing that although n-back WM performance seems to be predicted better, the difference between “same task” and “similar task” score prediction is not significant.

A possible reason for the lack of support of our hypothesis might lie in the nature of the task used in the scanner. That is, a lot of paradigms were developed in experimental contexts (Hedge et al., 2018) and therefore optimized for inducing a robust effect across participants instead of assessing interindividual differences. This might especially be the case for experimental tasks used in the scanner. For example, the emotional face matching task (Hariri et al., 2002) used for EMO assessment was developed to induce robust amygdala activation, rather than capturing individual emotion processing abilities. Additionally, the tasks used here were rather short and may have lacked enough difficult items for a clearer differentiation between participants. The n-back task, for example, most strongly differentiates between individuals when using high-load conditions (>3-back), both in terms of behavior and brain activity (Lamichhane et al., 2020). Therefore, using behavioral measurements from tasks optimized for obtaining stable group-average effects might have counteracted the successful prediction of interindividual differences.

### Is there network specificity for brain–behavior predictions?

We based the network specificity hypothesis on the assumption that if networks are reliably engaged during a task, then these networks should play an important role in the task outcome (i.e. specific performance). Importantly, our aim to demonstrate network specificity was based on the idea that *a priori* task-defined networks improve interpretability (Bzdok et al., 2012; Langner et al., 2018; Müller et al., 2018; Rottschy et al., 2012) as they reflect interactions between regions that are jointly engaged during a specific task and should therefore be biologically meaningful (J. Chen et al., 2021; Nostro et al., 2018; Pläschke et al., 2017). However, our results showed that prediction performance was weak regardless of the networks used (see Figure 3– COD). Comparison of the differences between networks showed that prediction from the whole-brain representation (Power et al., 2011) significantly improved prediction compared to the task specific networks (see Figure 4 D-F). The reason for an advantage of the whole-brain connectome remains to be revealed. We assume that some subtle pieces of information in the whole-brain connectome, which are not captured by the task networks, reflect individual processing differences in some parts of the tasks at hand and thus contribute to some extent to behavior prediction. Additionally, the whole-brain connectome has significantly more nodes than the task networks studied here, giving the model much more features to learn from. Hence, our results suggest that there is no network specificity, which is in line with the findings of Heckner et al., 2023 and Pläschke et al., 2017, 2020. Using networks based on group analyses may therefore not be a suitable avenue for assessing individual differences (Finn et al., 2017; O’Connor et al., 2017; Shah et al., 2016). Brain mapping results from group analyses typically reveal regions with low inter-subject variability (Hedge et al., 2018), and group-averaged patterns of brain activation often look quite different from patterns observed on the individual subject level (Miller et al., 2002). In addition, it has been shown that brain regions, for which activation has been found to be associated with behavioral outcomes, are not necessarily those that show up in standard group-average analyses (Ganis et al., 2005). Our results now indicate that this might also apply to networks derived from large samples that also reflect small average effects (HCP-derived networks) or from large-scale meta-analyses. However, improvement may be gained through an individualization of the networks prior to prediction. For example, Kong and colleagues employed a multi-session hierarchical Bayesian model to estimate individual-specific cortical network parcellations, significantly improving prediction performance relative to other parcellations (Kong et al., 2019). Similarly, using a different approach (Li et al., 2019) demonstrated an improvement in prediction accuracy using an iterative search based on a population-based functional atlas in combination with a map of inter-individual variability (D. Wang et al., 2015).

### Methodological considerations

In this study we aimed to predict complex behavior based on functional connectivity. Generally, our prediction accuracies were rather low. Nevertheless, they are comparable to the accuracies (correlation between predicted and observed score) reported in the literature (Dubois, Galdi, Han, et al., 2018; Greene et al., 2018; Heckner et al., 2023; Kandaleft et al., 2022; Ooi et al., 2022; Tomasi & Volkow, 2020). However, using correlation alone as a measure for prediction accuracy can skew the picture. All measures individually (correlation, COD, RMSE) have been shown to have their drawbacks and therefore it has been suggested that they should be considered together as a whole (Poldrack et al., 2020). Our results emphasize the importance of using more than one measure and especially using more than Pearson’s r as a measure for prediction performance, as this metric, when used in isolation, may draw an overly optimistic picture. As illustrated in our plots (see supplementary material), Pearson’s r invites the observer to interpret some apparent patterns. Yet, when looking at the model fit given by COD values, it can be easily seen that most models barely fit the data (see e.g. Figure 3). Surprisingly, only few prediction studies in the neuroimaging literature have reported metrics other than r. However, if they did, results were rather similar to ours, with finding only small amounts of variance explained (COD) and reporting high prediction errors on average (Dubois, Galdi, Han, et al., 2018; Kandaleft et al., 2022; Ooi et al., 2022).

There may be several reasons why we did not successfully predict behavioral performance. One is predicting behavioral scores of single tasks or questionnaires, like WM, as opposed to compound scores across many tests, like overall cognition (Akshoomoff et al., 2013; Dubois, Galdi, Paul, et al., 2018). Studies using compound scores generally report better accuracies (McCormick et al., 2022; Ooi et al., 2022), as they may capture individual abilities better and show higher reliabilities compared to individual test scores (Hedge et al., 2018). However, the interpretation and biological foundation of compound scores is debatable (Dubois, Galdi, Paul, et al., 2018; McFarland, 2012; Van Der Maas et al., 2006). In this study, we aimed to investigate specificity, and hence we focused on individual tasks or questionnaires at the cost of a potential decrease in prediction performance.

Another reason for the low prediction performance, related to the first explanation, might be the reliability of the predicted measures but also the features, setting an upper bound for detecting relationships (Cohen et al., 2013; Vul et al., 2009; Yarkoni & Braver, 2010). Using the HCP test– retest sample calculation of the correlations between measurement time points 1 and 2 (test–retest reliability) of the scores we used revealed reliabilities between 0.5 and 0.8, with highest reliabilities for the WM domain. In our and other studies, WM or intelligence scores were generally predicted better than other cognitive measures (Avery et al., 2020; Kandaleft et al., 2022; Ooi et al., 2022; Sripada et al., 2020; Takeuchi et al., 2021), which could be because these constructs are measured more accurately than others.

Finally, for ML applications in CV schemes, sample size is an important factor for achieving good prediction performance. The more data is available, the better a model can learn. In our case, our sample size decreased due to our carving out a subsample for *a priori* network delineation, leaving us with 420 subjects in the training set. This step was essential to assess network specificity using networks as close as possible to the investigated tasks. Other studies using the HCP dataset and similar algorithms have in part achieved slightly better predictions, possibly through larger training sets (Jiang et al., 2020; Ooi et al., 2022). The effects we sought to detect are presumably very small, hence a substantially larger dataset and/or more reliable behavioral assessments could be required to detect them (Marek et al., 2022).

### Limitations and outlook

We are aware that there is a plethora of preprocessing pipelines and feature selection models that may improve prediction. We used a well-established preprocessing pipeline (Glasser et al., 2016) and widely used ML models that previously yielded the highest predictions (Greene et al., 2018; Jiang et al., 2020; Yeung et al., 2022). Given that we saw a similar pattern of prediction accuracies irrespective of the model used, we would not expect a substantial change of the result pattern if other models were used.

The HCP dataset comprises young and healthy adults, with an above-population average intelligence. As the majority of subjects in the HCP dataset were highly educated, performed generally well on the tests, and as tests are optimized for group effects, the between-subject variability in this dataset is relatively limited, that is, suboptimal for approaches relying on individual differences. Nevertheless, the HCP currently offers the only dataset that allows for the investigation of such complex research questions as state specificity, state–target similarity, and network specificity in brain-based prediction settings, because it covers a vast array of phenotypic domains, both in and outside the scanner, while providing high-quality fMRI data in task and resting states in a large number of participants.

Finally, we aimed to cover a broad range of task network representations by using both ALE meta-analyses based on previously published neuroimaging results (see supplemental material), as well as networks from a high-powered single study using task fMRI data. For comparison with the task networks we included a whole-brain representation using the nodes from (Power et al., 2011). We acknowledge that different whole-brain representations, such as the parcellation by Schaefer and colleagues (Schaefer et al., 2018), could yield different and possibly even better prediction accuracies. We here, however, limited our analyses to one whole-brain representation, as our focus was on task networks and their interpretability.

## Conclusions

Here, using state-of-the-art ML algorithms for out-of-sample prediction analyses, we aimed to investigate the specific influence of the factors state, task, and network on behavior prediction from FC patterns. Based on previous research on brain–behavior relationships, we hypothesized that FC features from corresponding state, tasks, and networks would be more informative than non-corresponding features and hence improve prediction. We only found improvement for using task over resting state fMRI, as well as better predictions for whole brain compared to task specific networks. However, across three behavioral domains, predictive performance was generally poor, and there were no significant patterns indicating specificity of state, networks, or task similarity, when looking at RMSE and COD. A significant improvement of prediction performance based on task-fMRI (vs. resting-state fMRI) was only observed for the WM domain. Of note, an isolated consideration of Pearson’s correlation coefficient as the sole index of model fit would have led us to different and apparently overly optimistic conclusions. Hence, even with maximum state–network– behavior compatibility, the relationship between FC and behavior remains low. This study therefore emphasizes the need for a critical assessment of prediction accuracies and suggests that individual behavior cannot be successfully predicted based solely on FC in task-specific networks.

## Data availability

The data used in this study was obtained from the Human Connectome Project (HCP) database, which is publicly accessible at https://www.humanconnectome.org/. However some parameters needed application for restricted access, which all authors handling the data had granted by the HCP. The three meta-analytically defined networks will be openly available via the ANIMA-database (Reid et al., 2016; https://anima.fz-juelich.de/).

## Acknowledgments

We would like to sincerely thank all participants and contributors to the data provided by the Human Connectome Project, WU-Minn Consortium (Principal Investigators: David Van Essen and Kamil Ugurbil; 1U54MH091657) funded by the 16 NIH Institutes and Centers that support the NIH Blueprint for Neuroscience Research; and by the McDonnell Center for Systems Neuroscience at Washington University.

The work was supported by: Helmholtz Portfolio Theme “Supercomputing and Modeling for the Human Brain”; European Union’s Horizon 2020 Research and Innovation Programme under Grant Agreement No. 945539 (HBP SGA3).

## Author contributions

Nevena Kraljević: Conceptualization, Methodology, Software, Formal analysis, Writing - Original Draft, Writing – review & editing, Visualization. Robert Langner: Conceptualization, Methodology, Supervision, Writing - Review & Editing. Vincent Küppers, Writing - Review & Editing: Methodology. Federico Raimondo: Methodology, Software, Writing - Review & Editing. Kaustubh R. Patil: Methodology, Software, Writing - Review & Editing. Simon B. Eickhoff: Funding acquisition, Resources, Supervision, Writing - Review & Editing. Veronika I. Müller: Conceptualization, Methodology, Supervision, Writing - Review & Editing.

## Corresponding authors

Nevena Kraljević, Veronika Müller.

