## Supplemental Material for "Network and State Specificity in Connectivity-Based Predictions of Individual Behavior"

Nevena Kraljević[1, 2], Robert Langner[1, 2], Vincent Küppers[1, 3], Federico Raimondo[1, 2], Kaustubh R. Patil[1, 2], Simon B. Eickhoff[1, 2], Veronika I. Müller[1, 2]

### Methods

Overview of references per task:

*Table S1: Behavioral Scores used in the prediction*

| Domain | "Same" / In-scanner task | "Similar " / Out-of-scanner task |
|---|---|---|
| Working Memory / WM | N-back (Barch et al., 2013) | List sorting (NIH Toolbox List Sorting Working Memory Test; (*Cognition Measures*, n.d.) |
| Theory of mind / SOCIAL | Labelling of interaction between animated shapes as random or interaction (Castelli et al., 2000; Wheatley et al., 2007) | Compound score "Social Satisfaction" (Babakhanyan et al., 2018) composed of scores for:<br>Friendship, loneliness, emotional support, instrumental support, and perceived rejection all from |

| | | |
|---|---|---|
| | | NIH Toolbox Emotion battery (*Emotion Measures*, n.d.; Salsman et al., 2013). |
| Emotion Recognition / EMO | Face-matching. Adapted by (Hariri et al., 2002) | Penn Emotion Recognition Test (Gur et al., 2002, 2010) |

### *Network Delineation*

#### Network Delineation via Meta-Analysis

As a second approach, to offer feature spaces that are entirely independent from the target, we performed three activation likelihood estimation (ALE) meta-analyses for each of the selected tasks: WM, SOCIAL, and EMO. WM and EMO were based on previous meta-analyses (WM: Rottschy et al., 2012;  EMO: Müller, Höhner, et al., 2018), but were extended by including recent publications (findings up to March 2020) and reduced to those tasks that matched the three tasks used in the HCP (i.e. only 2-back vs. 0 back experiments for WM, matching faces > matching shapes for EMO). For SOCIAL we performed our own literature search and coding procedure, following the guidelines for neuroimaging meta-analyses (Müller, Cieslik, et al., 2018) and including experiments that used a theory of mind task using animated shapes and report results of the interaction > random contrast. For each specific task, a meta-analysis was calculated using the ALE algorithm (details about the method see Müller, Cieslik, et al., 2018 and Kogler et al., 2020. From these resulting ALE maps, illustrating spatial convergence across experiments, we extracted all peak coordinates with a minimum distance of 15 mm using FSL. This resulted in three networks from the meta-analyses: MetaWM, MetaSOCIAL, and MetaEMO. The three meta-analytically defined networks will be openly available via the ANIMA-database (Reid et al., 2016; https://anima.fz-juelich.de/).

### *Network-based prediction of individual behavior*

In addition to PLS, we used Support Vector Regression (SVR), Random Forest, kernel ridge regression algorithms for prediction. Lastly, we performed connectivity-based prediction modelling (CBPM; Finn et al., 2015; Shen et al., 2017) as a popular feature reduction technique with both PLS and kernel ridge as algorithms.

For each algorithm we tuned the hyperparameters in an inner 5x-CV loop. For SVR we ran two different kernels: linear and RBF-kernel. For the linear kernel we tuned the regularization parameter C within [1e-6, 1e-5, 1e-4, 0.0005, 0.001, 0.005], with maximum 2000 iterations within solver. For the RBF-kernel we used the same regularization parameter range, but extended it by [0.01, 0.1, 1, 5, 10]. For the random forest prediction, the number of trees was set to 2000, with mean squared error as the criterion. The number of features was tuned within [0.14, 0.22, 0.33, 0.5, 0.75], with a minimum number of 5 samples required to be at a leaf node. For kernel ridge regression we tuned the lambdas in a range from 0-1000000. For the connectivity-based prediction modelling (Finn et al., 2015; Shen et al., 2017) we used the same hyperparameter tuning as without the feature reduction method.

For the significance tests, we used the Nadeau-Bengio machine learning adjusted t-test (Nadeau & Bengio, 1999): $t = \frac{\frac{1}{n}\sum_{j=1}^{n} x_j}{\sqrt{(\frac{1}{n} + \frac{n_{test}}{n_{train}})\hat{\sigma}^2}}$. Within each cognitive domain, we first tested effects of state and network by averaging prediction performance of the respective other factors (i.e., averaging across networks and task when testing for state effects, and across state and task when testing for network effects). As domain specificity is an extension of state specificity, we here only averaged across networks for same and similar tasks, respectively. Significant effects (corrected for multiple comparisons) were then further assessed by comparing the respective individual prediction scores between each other. In particular, to assess i) state specificity we compared the prediction performance between states, while keeping network and task constant. That is, we only compared predictions between corresponding networks and tasks (e.g. comparing the prediction performance of “same” WM task score based on FC within Power nodes in resting state to the prediction performance of “same” WM task score based on FC within Power nodes in WM state). To assess ii) network specificity we compared the prediction performance between all networks, while keeping state and task constant (e.g. comparison of prediction performance of “same” WM score in resting state WM networks compared to “same” WM score in resting state in EMO networks). To assess iii) domain specificity we compared the prediction performance of “same” and “similar” task scores, while keeping state and network constant (e.g. comparison of prediction performance of “same” WM task score based on Power nodes in WM state to prediction performance of “similar” WM task score based on Power nodes in WM state).

### Results

*Table S2. List of coordinates for WM-NW and WM-Meta.*

| MNI coordinates peak voxel WM-NW | | |
|---|---|---|
| X | Y | Z |
| 34 | -58 | -32 |
| 46 | -46 | 46 |
| -30 | -60 | -32 |
| -6 | 18 | 48 |
| 32 | 6 | 58 |
| -28 | 6 | 54 |
| -44 | -52 | 46 |
| 40 | 34 | 28 |
| -8 | -64 | 50 |
| 8 | -68 | 54 |
| 34 | 22 | 4 |
| -32 | 50 | 16 |
| -32 | 20 | 0 |
| -44 | 26 | 34 |
| 38 | -60 | -48 |
| 12 | -76 | -24 |
| 38 | 48 | 18 |
| -8 | -80 | -26 |
| -16 | 8 | 12 |
| 18 | 10 | 16 |
| 58 | -30 | -14 |
| -8 | -58 | -54 |
| 52 | 10 | 16 |
| 24 | 46 | -14 |
| -12 | -92 | 2 |
| 10 | 2 | 6 |
| 0 | -50 | -18 |
| 0 | -30 | -4 |
| 2 | -18 | -18 |
| 8 | -58 | -54 |
| 48 | 6 | 30 |
| -24 | 50 | -12 |
| 2 | -12 | 16 |
| 0 | -62 | -36 |
| -56 | -36 | -14 |
| 2 | 12 | 24 |

| MNI coordinates peak voxel WM-Meta-NW | | |
|---|---|---|
| X | Y | Z |
| -46 | 6 | 36 |
| -28 | 2 | 54 |
| -46 | 26 | 28 |
| -34 | -54 | 48 |
| 42 | -46 | 44 |
| 10 | -66 | 52 |
| -2 | 18 | 48 |
| -2 | 32 | 38 |
| 30 | 8 | 56 |
| 32 | 24 | -2 |
| 44 | 34 | 26 |
| -32 | -60 | -34 |
| 30 | -60 | -30 |
| -32 | 22 | 0 |
| -38 | 50 | 8 |
| 10 | -76 | -24 |
| -8 | -76 | -28 |
| -12 | -68 | 60 |
| -16 | -2 | 16 |

| | | |
|---|---|---|
| 0 | -36 | -42 |
| -2 | -32 | 24 |
| -44 | -50 | 20 |
| 2 | -22 | -36 |
| 28 | -58 | 66 |
| -14 | -26 | -32 |
| 20 | -28 | 14 |
| 20 | -96 | -14 |
| 34 | -90 | -18 |
| 16 | 28 | -22 |
| -18 | -46 | 24 |
| 24 | -20 | -6 |
| -22 | -58 | 0 |

*Table S3. List of coordinates for SOCIAL-NW and SOCIAL-Meta.*

| MNI coordinates peak voxel SOCIAL-NW | | |
|---|---|---|
| **X** | **Y** | **Z** |
| -12 | -94 | 18 |
| -22 | -52 | 64 |
| 22 | -50 | 68 |
| 18 | -86 | 24 |
| -10 | -80 | 34 |
| -18 | -12 | 70 |
| 10 | -80 | -6 |
| -26 | -44 | 12 |
| 12 | -74 | 38 |
| 16 | -10 | 74 |
| 26 | -44 | 10 |
| 2 | -24 | 30 |
| 28 | 66 | 2 |
| 48 | -56 | 46 |
| 40 | 54 | -6 |
| -26 | 68 | 4 |
| 0 | 16 | 10 |
| 40 | 48 | 10 |
| 50 | -2 | -2 |
| -20 | -26 | 76 |
| -2 | -10 | 70 |
| 2 | 40 | 16 |
| -46 | -10 | 60 |
| -22 | -76 | 4 |
| 42 | 38 | 32 |
| -12 | 26 | 6 |
| -48 | -2 | -4 |

| MNI coordinates peak voxel SOCIAL-Meta-NW | | |
|---|---|---|
| **X** | **Y** | **Z** |
| 58 | -48 | 14 |
| -58 | -46 | 16 |
| 62 | -8 | -16 |
| 54 | 6 | -22 |
| 10 | 62 | 22 |
| -44 | -58 | -10 |
| 54 | 28 | 6 |
| -54 | 26 | 10 |
| 8 | -48 | 50 |
| 8 | -54 | 36 |
| -60 | -8 | -14 |

| | | |
|---|---|---|
| -8 | -76 | -4 |
| 38 | 14 | 8 |
| -14 | -28 | 24 |
| 4 | 46 | 2 |
| -42 | -26 | 62 |
| 48 | -8 | 52 |
| 20 | -26 | 76 |
| -62 | 2 | 20 |
| -46 | -36 | 16 |
| 2 | 4 | 50 |
| 14 | 26 | 8 |
| 42 | -22 | 62 |
| 20 | -30 | 24 |
| 2 | -30 | 50 |
| -42 | -60 | 48 |
| -8 | -10 | 52 |
| -26 | 46 | -12 |
| 6 | -2 | 22 |
| 66 | -20 | 8 |
| -42 | 12 | -8 |
| 14 | -16 | 46 |
| -38 | -14 | 24 |
| 60 | -2 | 38 |
| -36 | 12 | 10 |
| -28 | 44 | 36 |
| 36 | -10 | 20 |
| 30 | 22 | 60 |
| -12 | 18 | -16 |
| -26 | -30 | 34 |
| 40 | -66 | 8 |
| 12 | 48 | -26 |
| -14 | 32 | -22 |
| 2 | -70 | -20 |
| 44 | -36 | 34 |
| 44 | -32 | 18 |
| -32 | -52 | 30 |
| -46 | 32 | 38 |
| -20 | 12 | 30 |
| 62 | -26 | -16 |
| -46 | -66 | -38 |
| -64 | -30 | -14 |
| -26 | -48 | -50 |
| -34 | -34 | -38 |
| 48 | -62 | -38 |

*Table S4. List of coordinates for EMO-NW and EMO-Meta.*

| MNI coordinates peak voxel EMO-NW | | |
|---|---|---|
| **X** | **Y** | **Z** |
| 24 | -96 | -4 |
| 42 | -48 | -20 |
| -20 | -94 | -12 |
| 38 | -72 | -14 |
| 18 | -4 | -16 |
| -18 | -4 | -18 |
| -34 | -86 | -12 |
| -40 | -54 | -20 |
| 44 | 18 | 24 |
| -4 | -82 | 2 |
| 14 | -32 | -2 |
| -8 | -76 | -38 |
| 34 | 34 | -14 |
| 32 | -6 | -38 |
| -40 | 18 | 26 |
| 48 | -64 | 18 |
| -10 | -32 | -2 |
| -34 | -10 | -32 |
| 14 | -70 | 10 |
| 50 | -42 | 14 |
| -24 | -24 | -8 |
| -36 | 30 | -16 |
| -2 | -2 | -16 |
| 20 | -38 | -44 |
| 0 | -52 | -36 |
| 22 | -52 | 4 |
| -20 | -36 | -44 |
| -50 | -72 | 18 |
| 50 | -10 | -12 |
| 62 | -52 | 12 |
| 60 | -42 | -4 |
| 10 | -78 | -38 |
| 2 | 52 | -14 |
| 46 | 2 | 54 |
| -52 | -46 | 12 |
| 34 | -56 | 44 |
| -44 | 40 | -2 |
| -60 | -40 | -8 |
| 4 | -60 | 40 |
| 0 | 4 | 28 |
| 36 | -74 | 24 |

| MNI coordinates peak voxel EMO-Meta-NW | | |
|---|---|---|
| **X** | **Y** | **Z** |
| 20 | -4 | -18 |
| 28 | -94 | -6 |
| -22 | -6 | -14 |
| -22 | -96 | -6 |
| 42 | 12 | 28 |
| -42 | -54 | -22 |
| 40 | -50 | -26 |
| -18 | -32 | -2 |
| -50 | -48 | 4 |
| -54 | 18 | 32 |

| | | |
|---|---|---|
| 4 | 58 | 32 |
| 2 | 38 | 50 |
| 52 | 28 | -6 |
| -30 | -68 | -48 |
| -52 | -10 | -12 |
| 22 | -68 | 24 |
| 6 | -12 | 6 |
| 24 | -32 | -20 |
| -40 | 2 | 54 |
| 28 | -24 | 60 |
| -50 | 12 | 46 |
| 22 | 0 | 6 |
| 34 | 20 | 54 |
| 8 | -96 | 26 |
| -4 | -22 | 58 |
| 16 | -26 | 72 |
| 2 | 4 | -2 |
| -18 | -92 | -28 |
| 8 | 6 | 12 |
| 0 | -86 | 36 |
| -14 | -26 | 72 |
| 6 | -40 | 68 |
| -64 | -6 | 28 |
| -12 | 10 | 4 |
| 66 | -2 | 28 |
| -50 | -10 | 32 |
| -50 | -28 | -2 |
| -38 | 18 | 58 |
| 18 | -12 | 20 |
| -38 | -18 | 40 |
| 12 | -10 | 78 |
| 2 | 12 | 70 |
| -12 | -40 | 66 |
| 30 | -22 | 12 |
| -6 | -32 | -32 |
| -14 | -2 | 16 |
| -18 | 30 | 60 |
| -30 | -80 | 26 |
| -58 | -12 | 48 |
| 34 | -68 | -50 |
| -60 | -10 | -36 |
| -50 | -68 | 46 |
| -32 | -60 | 44 |

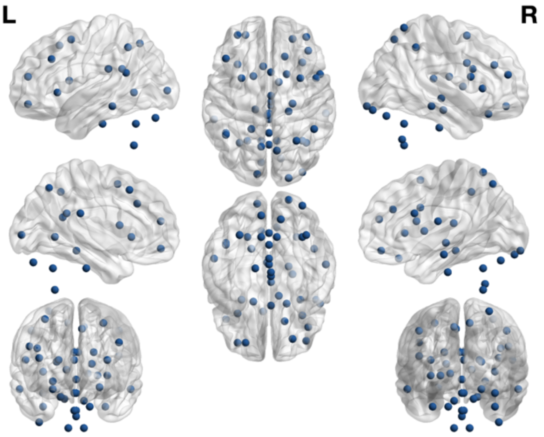


*Figure S1. WM Network Nodes.*

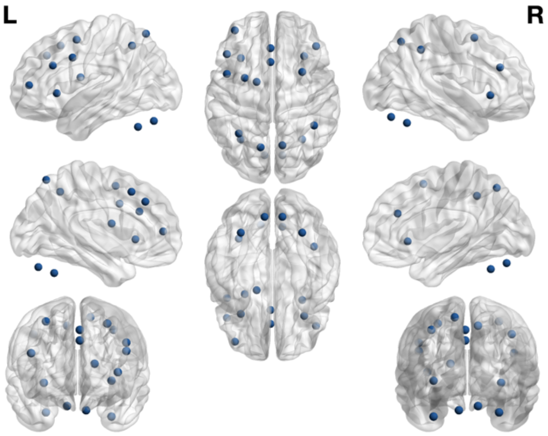


*Figure S2. WM Meta-Analysis Network Nodes.*

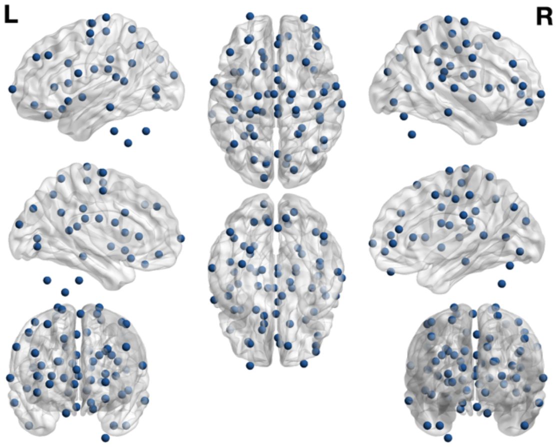


*Figure S3. SOCIAL Network Nodes.*

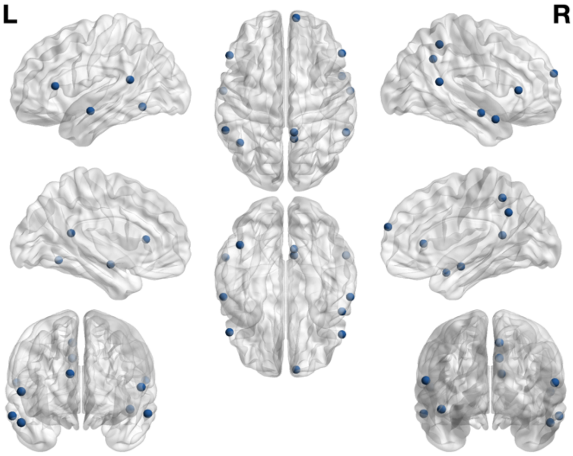


*Figure S4. SOCIAL Meta-Analysis Network Nodes*

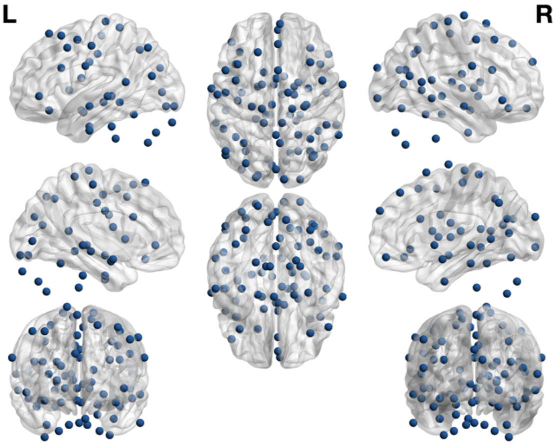


*Figure S5. EMO Network Nodes*

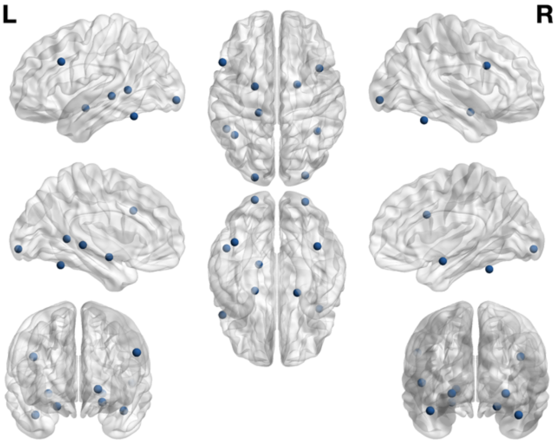


*Figure S6. EMO Meta-Analysis Network Nodes*

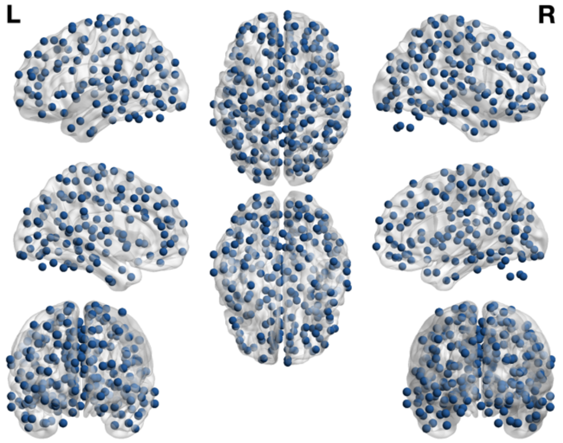

*Figure S7. Whole-brain Power nodes.*

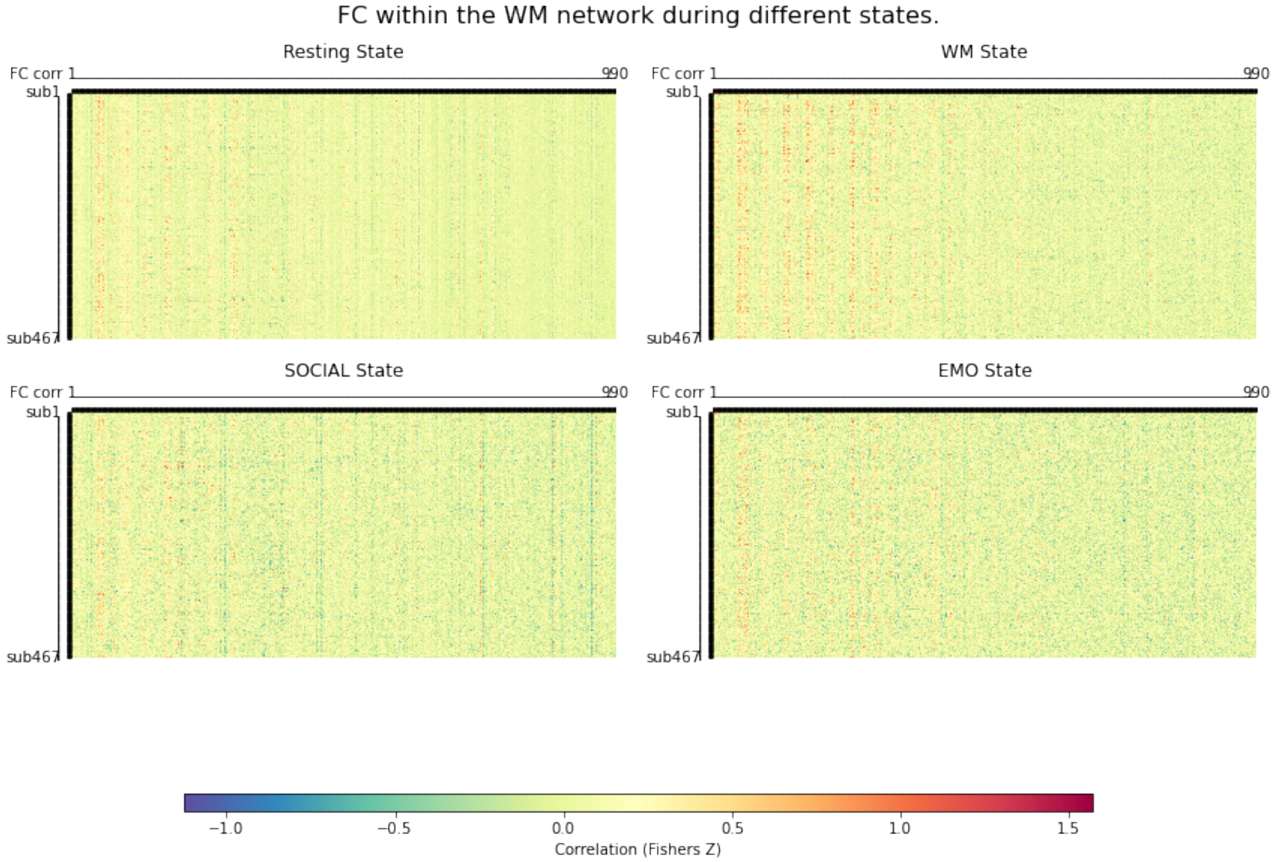


*Figure S8) Heatmap of FC within the WM network for all participants. FC reflects the Fisher Z- transformed Pearson correlation coefficients between all network nodes*

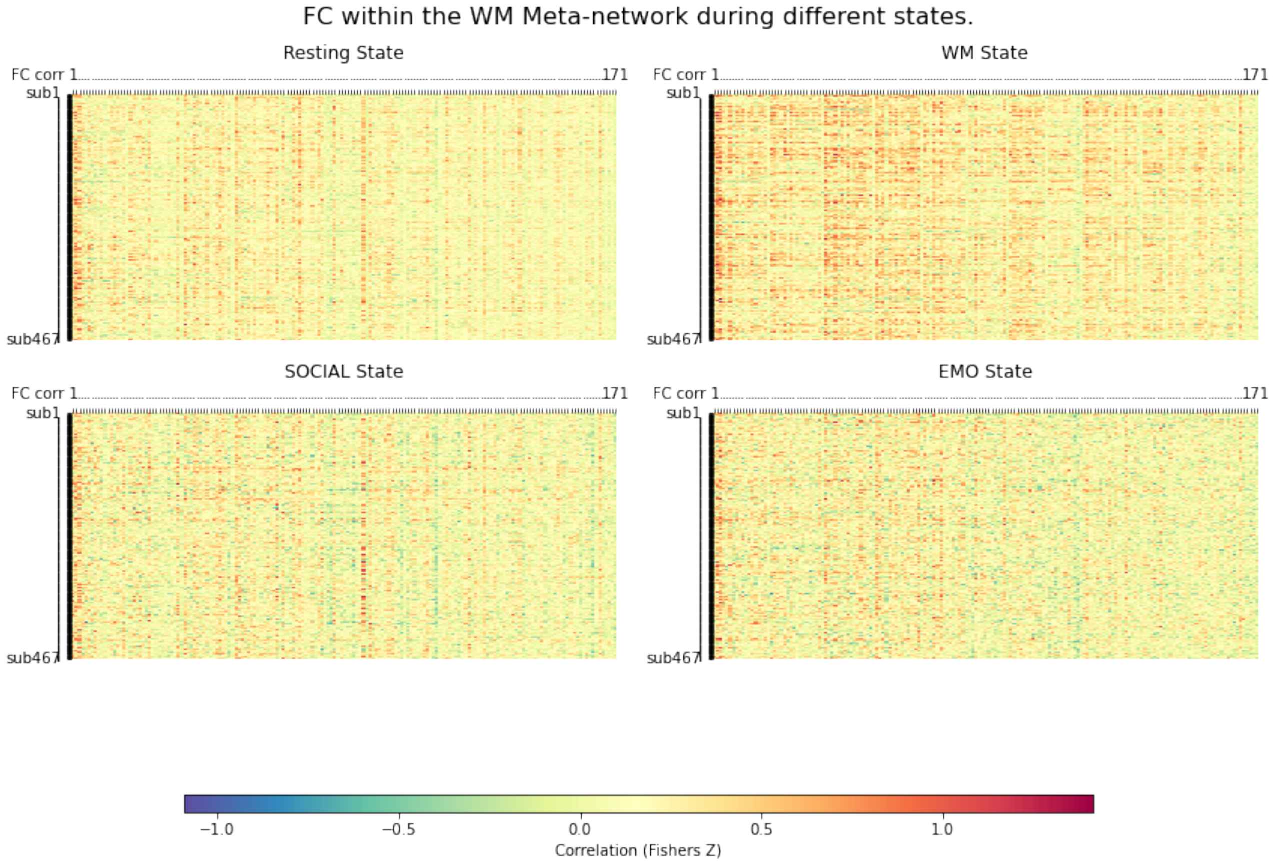


*Figure S9) Heatmap of FC within the WM-meta network for all participants. FC reflects the Fisher Z- transformed Pearson correlation coefficients between all network nodes.*

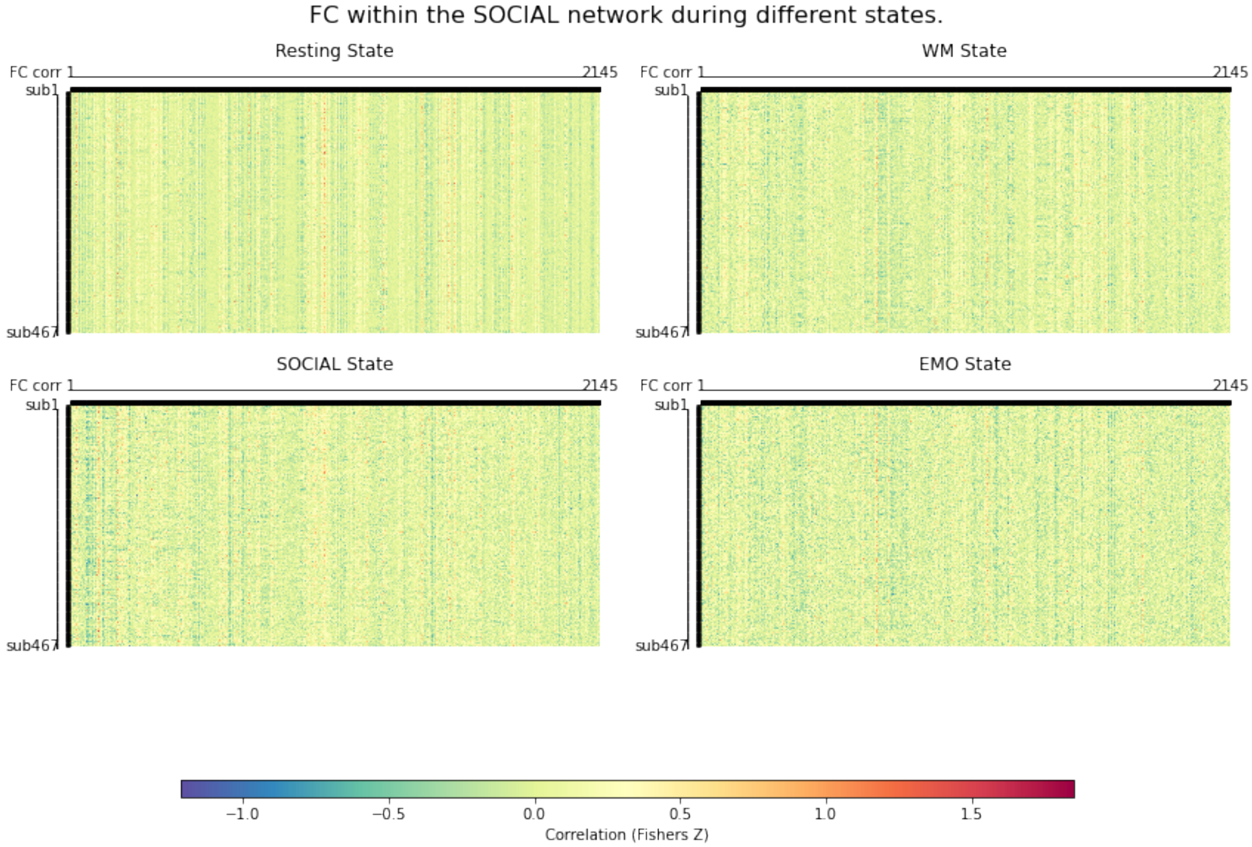


*Figure S10) Heatmap of FC within the SOCIAL network for all participants. FC reflects the Fisher Z- transformed Pearson correlation coefficients between all network nodes.*

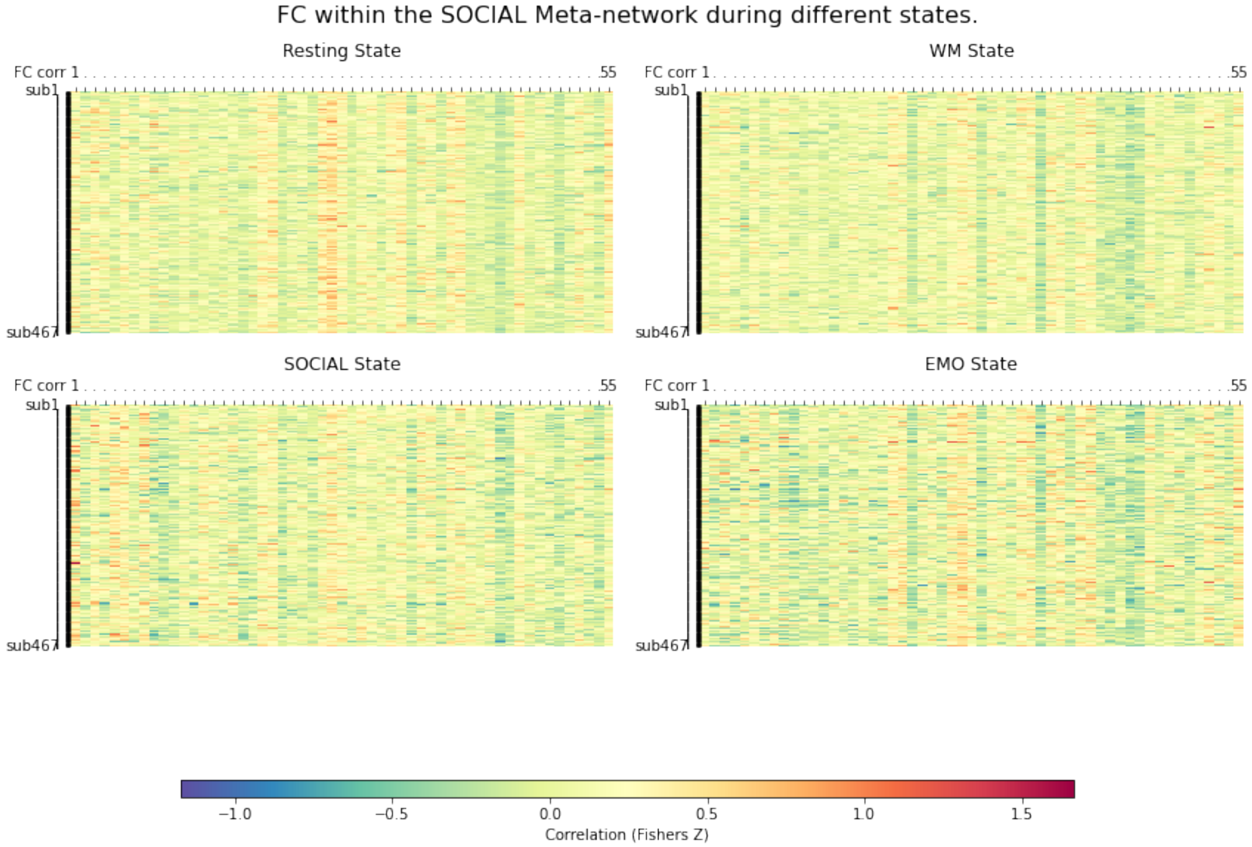


*Figure S11) Heatmap of FC within the SOCIAL-meta network for all participants. FC reflects the Fisher Z- transformed Pearson correlation coefficients between all network nodes.*

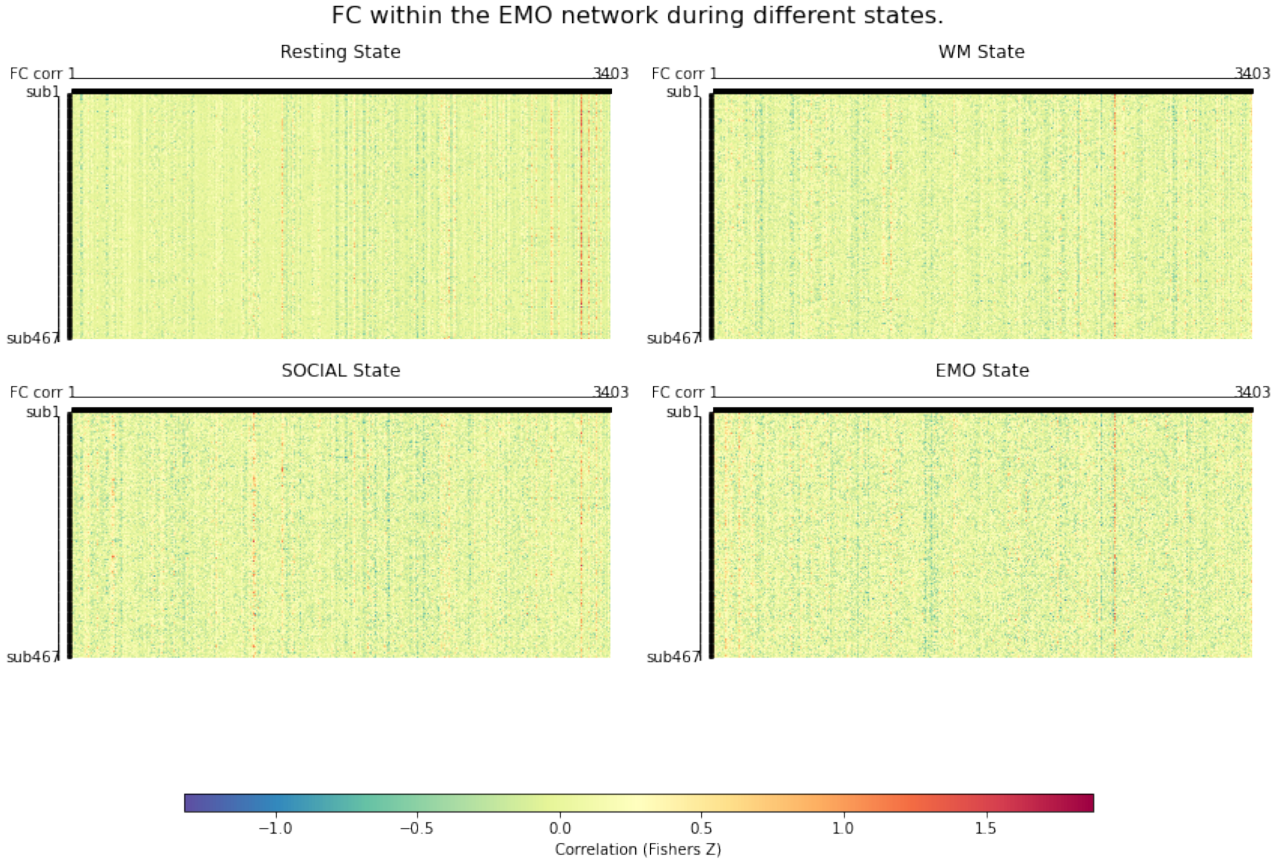


*Figure S12) Heatmap of FC within the EMO network for all participants. FC reflects the Fisher Z- transformed Pearson correlation coefficients between all network nodes.*

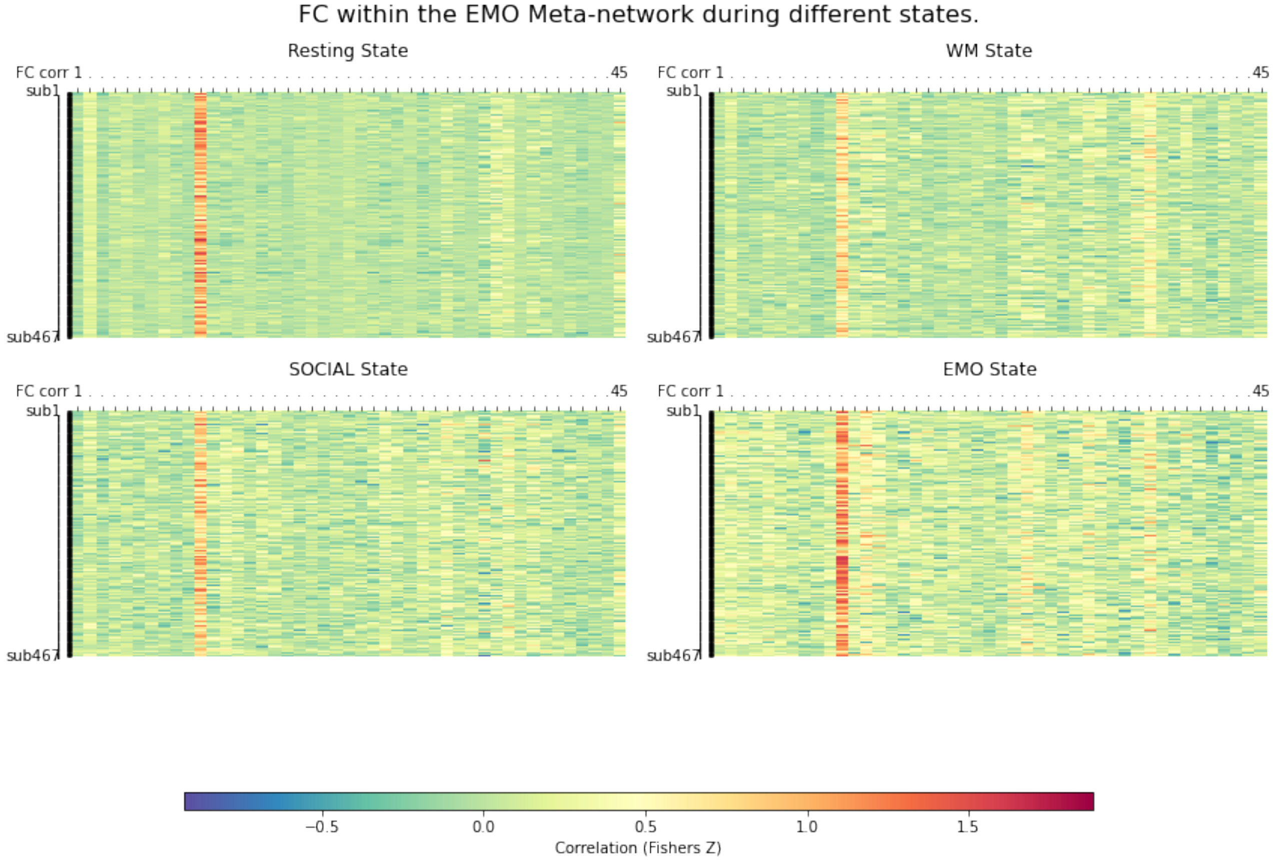


*Figure S13) Heatmap of FC within the EMO-meta network for all participants. FC reflects the Fisher Z- transformed Pearson correlation coefficients between all network nodes.*

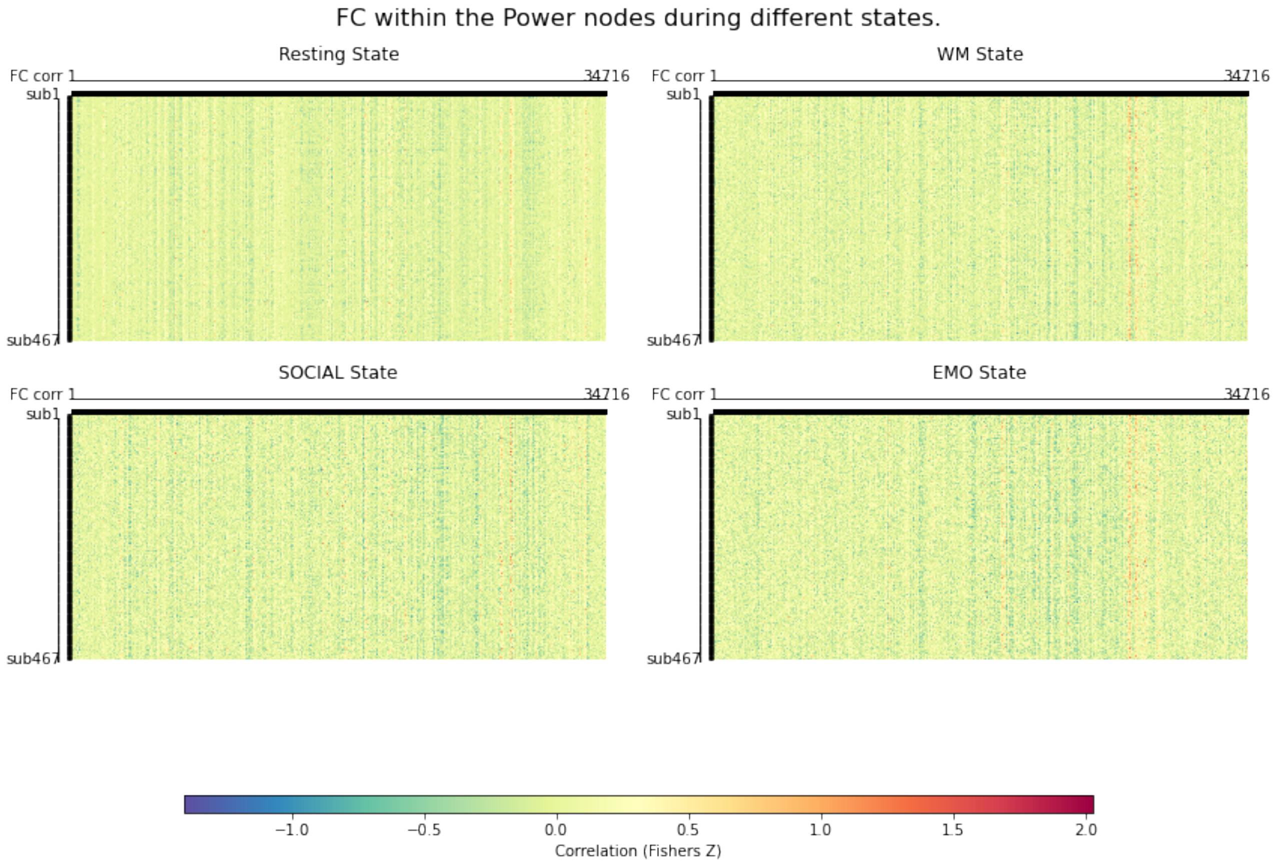


*Figure S14) Heatmap of FC within the Power nodes for all participants. FC reflects the Fisher Z- transformed Pearson correlation coefficients between all network nodes.*

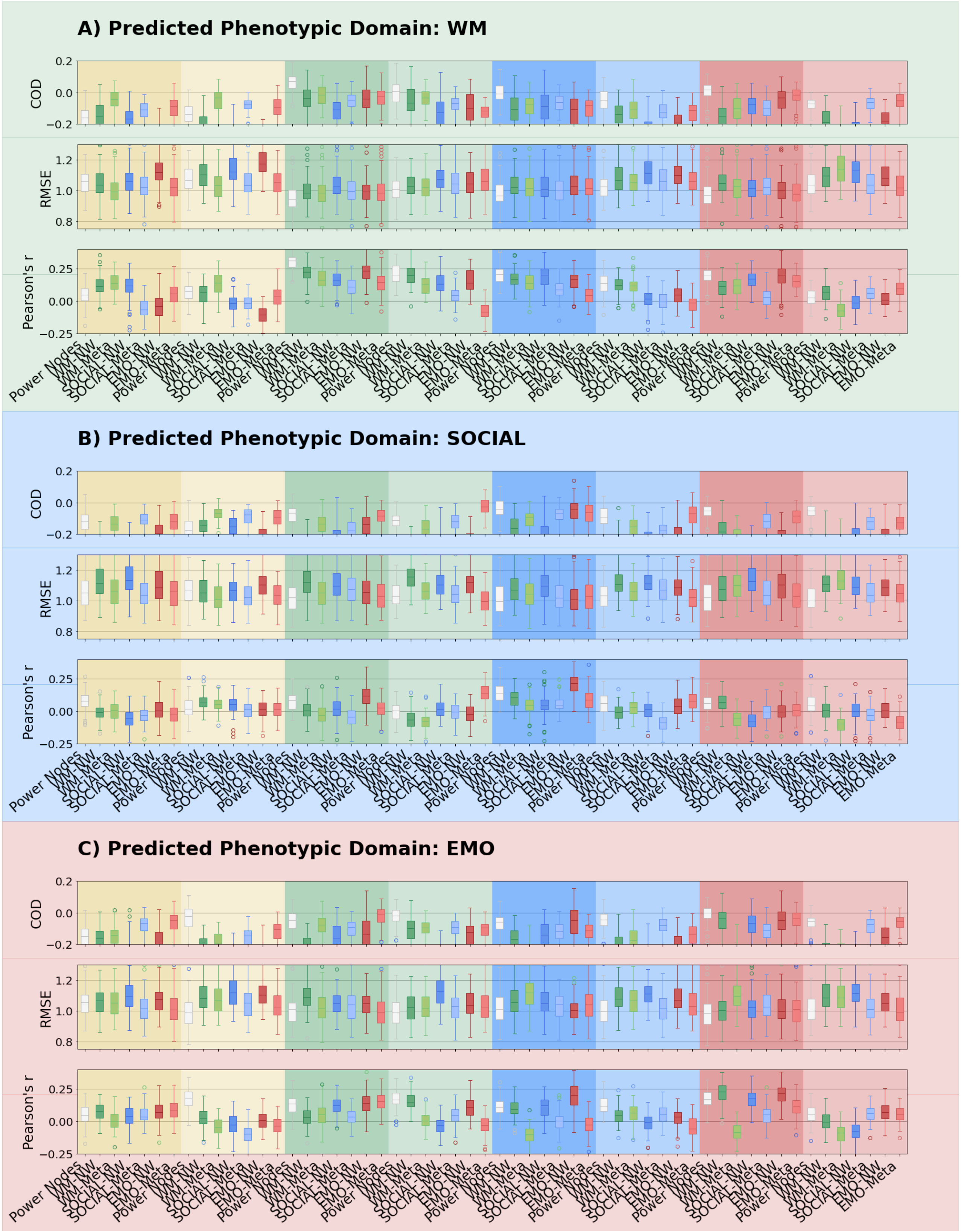


*Figure S15) PLS 100 x leave-30%-out CV*

Boxplots of the distribution of prediction accuracies from PLS 100 x leave-30%-out CV for WM, SOCIAL, and EMO domain, for coefficient of determination (COD) / model fit, RMSE and Pearson's r.

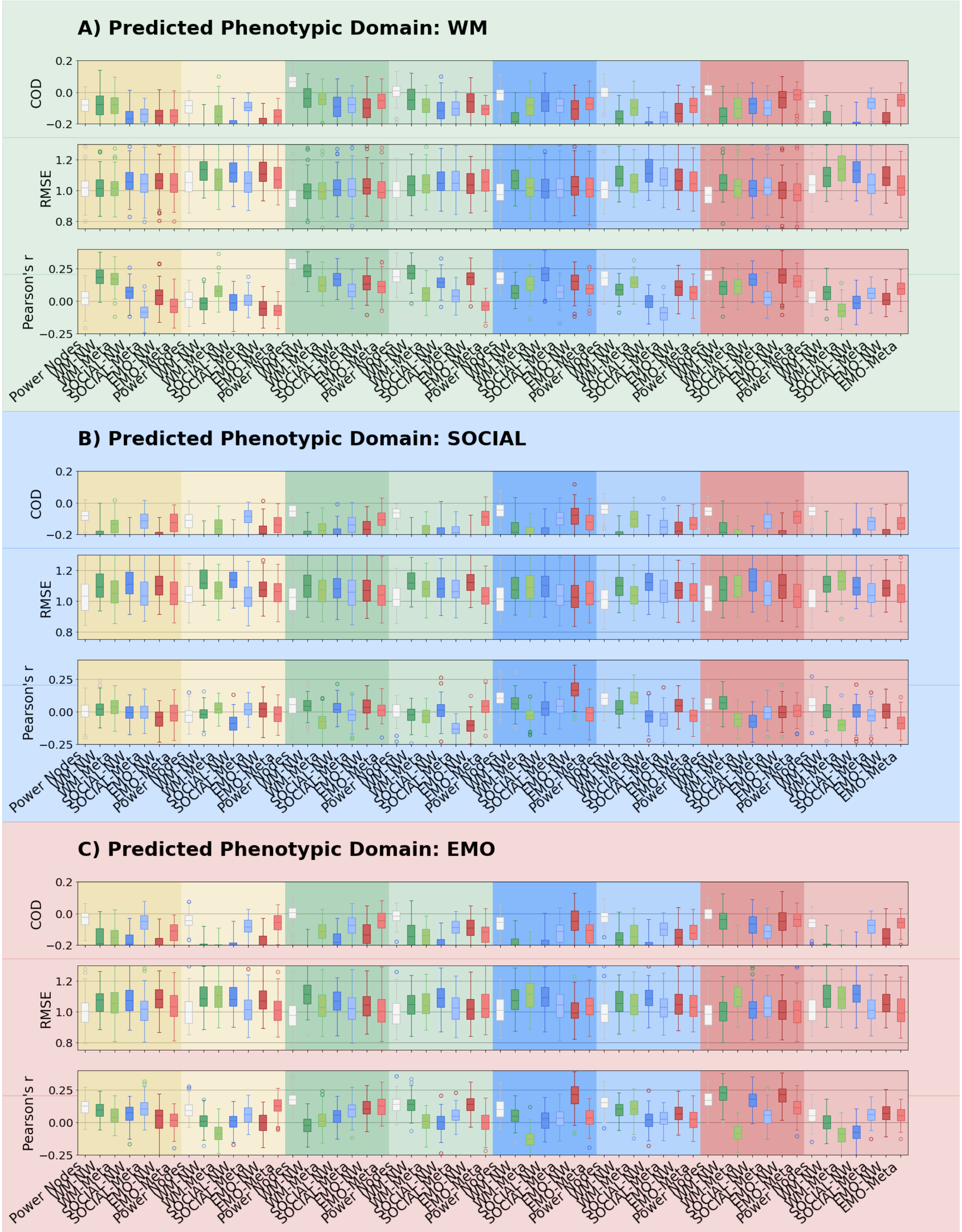


*Figure S16) PLS 100 x leave-30%-out CV – trimmed time series*

Boxplots of the distribution of prediction accuracies from PLS 100 x leave-30%-out CV – trimmed time series for WM, SOCIAL, and EMO domain, for coefficient of determination (COD) / model fit, RMSE and Pearson's r.

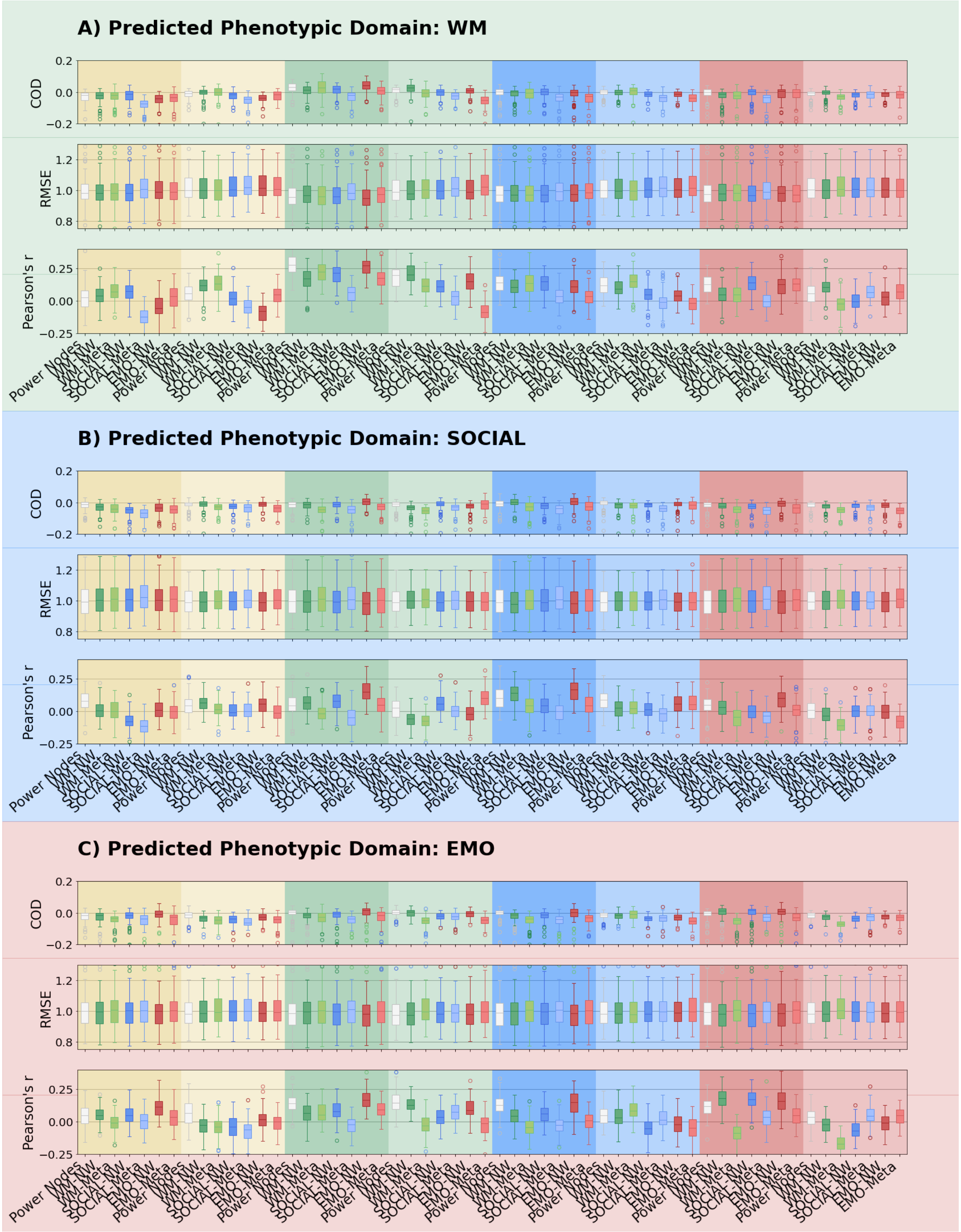

*Figure S17) Random Forest 100 x leave-30%-out CV*

Boxplots of the distribution of prediction accuracies from Random Forest 100 x leave-30%-out CV for WM, SOCIAL, and EMO domain, for coefficient of determination (COD) / model fit, RMSE and Pearson’s r.

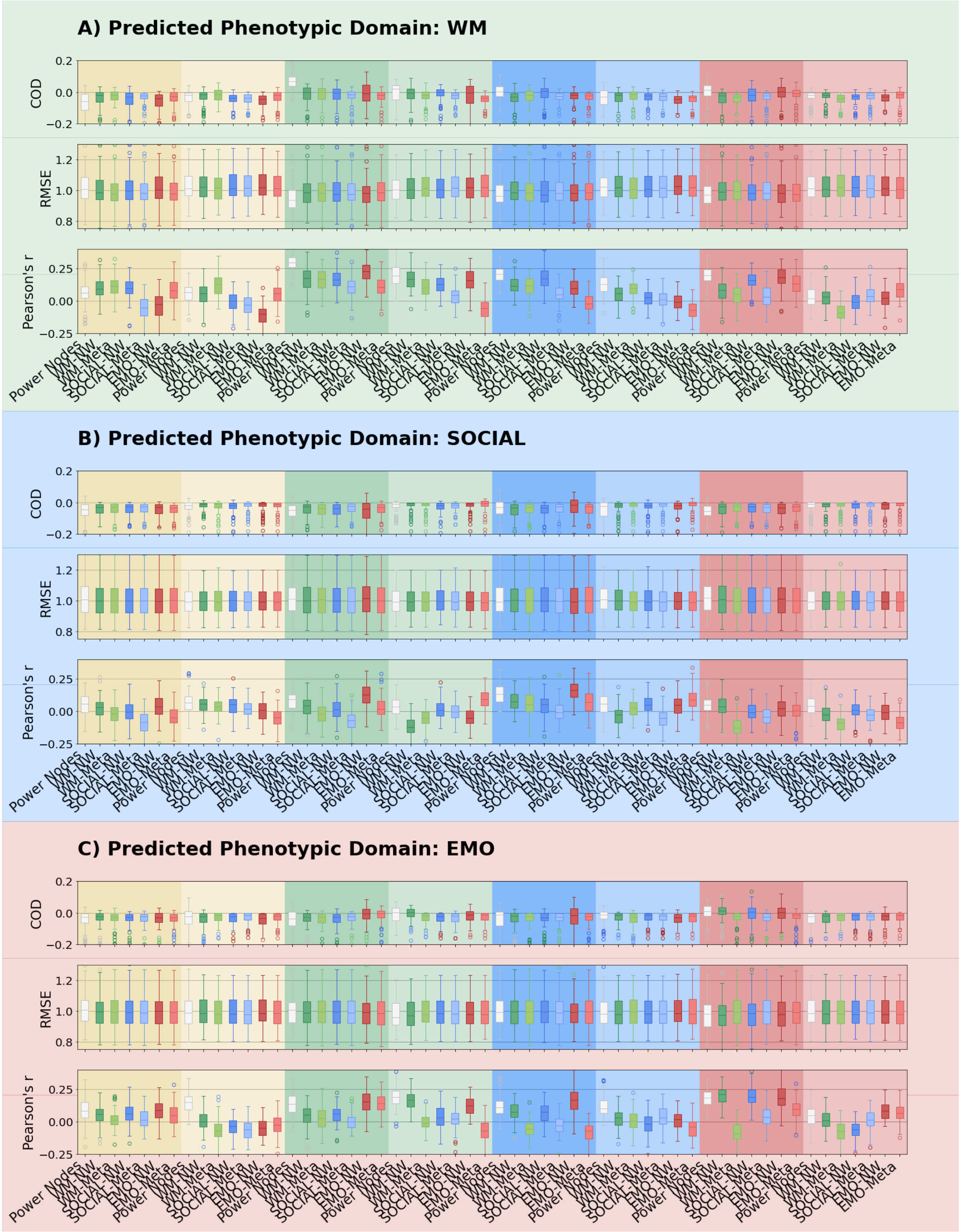


*Figure S18) SVR – linear kernel - 100 x leave-30%-out CV*

Boxplots of the distribution of prediction accuracies from SVR – linear kernel - 100 x leave-30%-out CV for WM, SOCIAL, and EMO domain, for coefficient of determination (COD) / model fit, RMSE and Pearson's r.

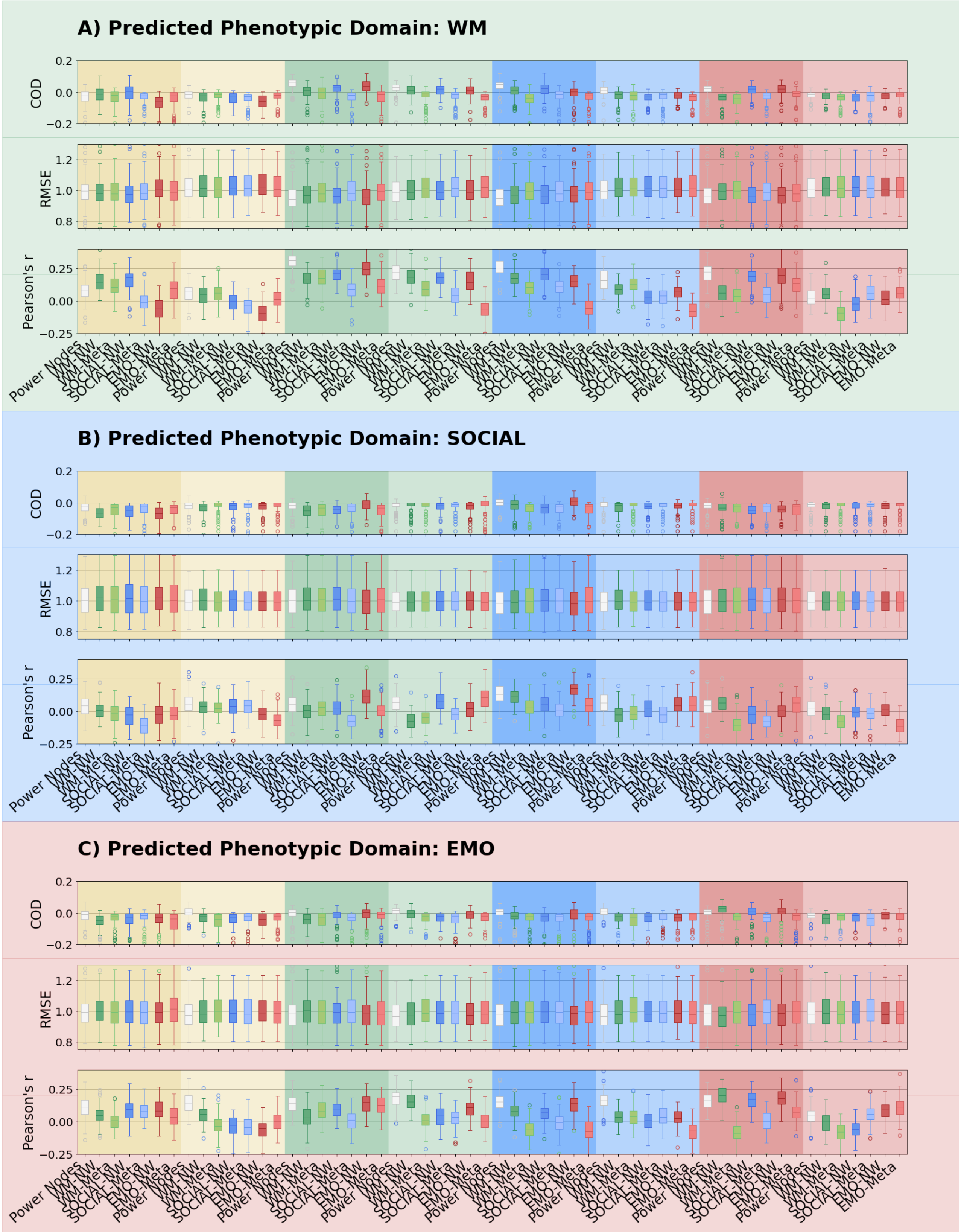


*Figure S19) SVR – rbf kernel - 100 x leave-30%-out CV*

Boxplots of the distribution of prediction accuracies from SVR – RBF kernel - 100 x leave-30%-out CV for WM, SOCIAL, and EMO domain, for coefficient of determination (COD) / model fit, RMSE and Pearson's r.

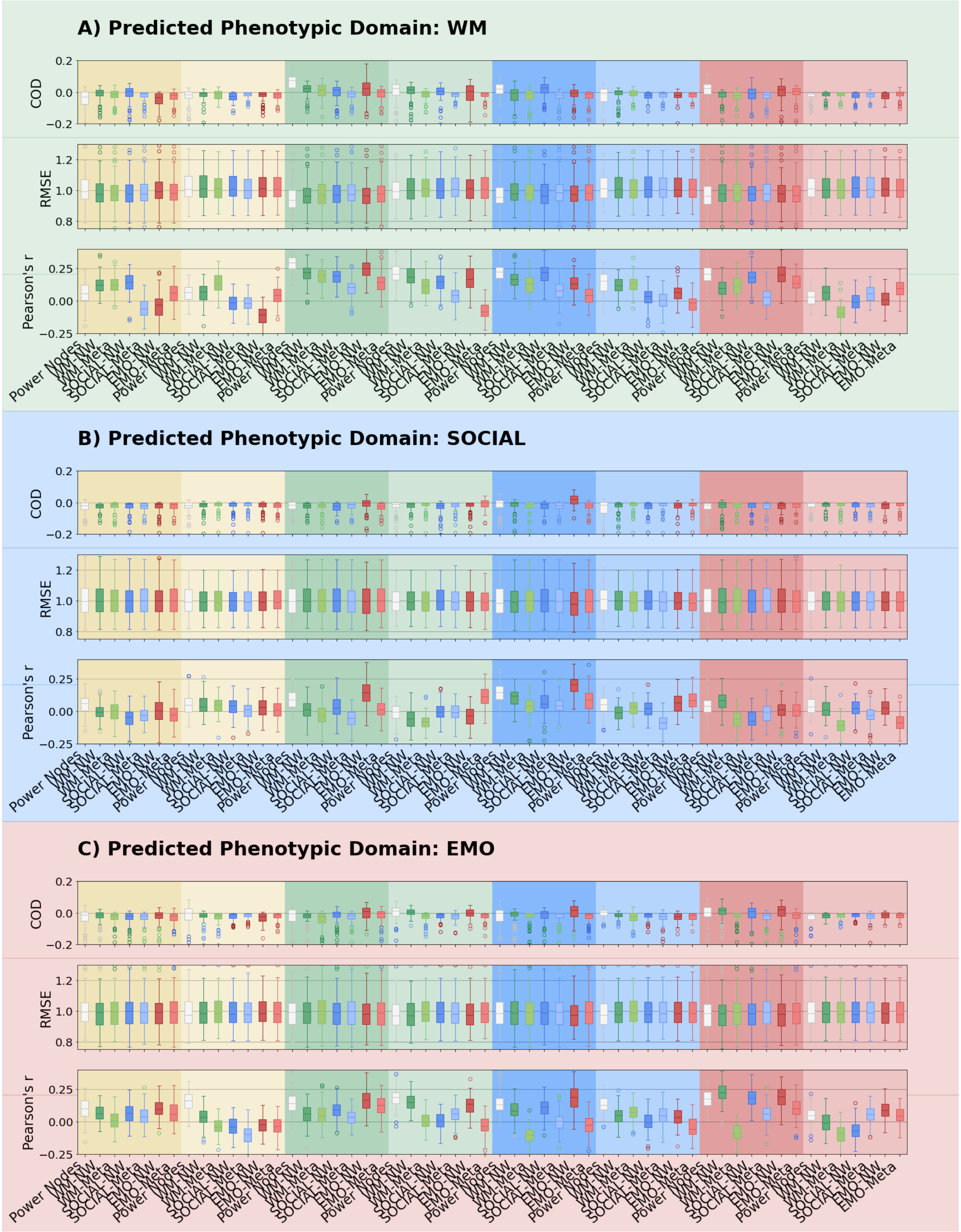


*Figure S20) Kernel Ridge Regression - 100 x leave-30%-out CV*

Boxplots of the distribution of prediction accuracies from Kernel Ridge Regression - 100 x leave-30%-out CV for WM, SOCIAL, and EMO domain, for coefficient of determination (COD) / model fit, RMSE and Pearson's r.

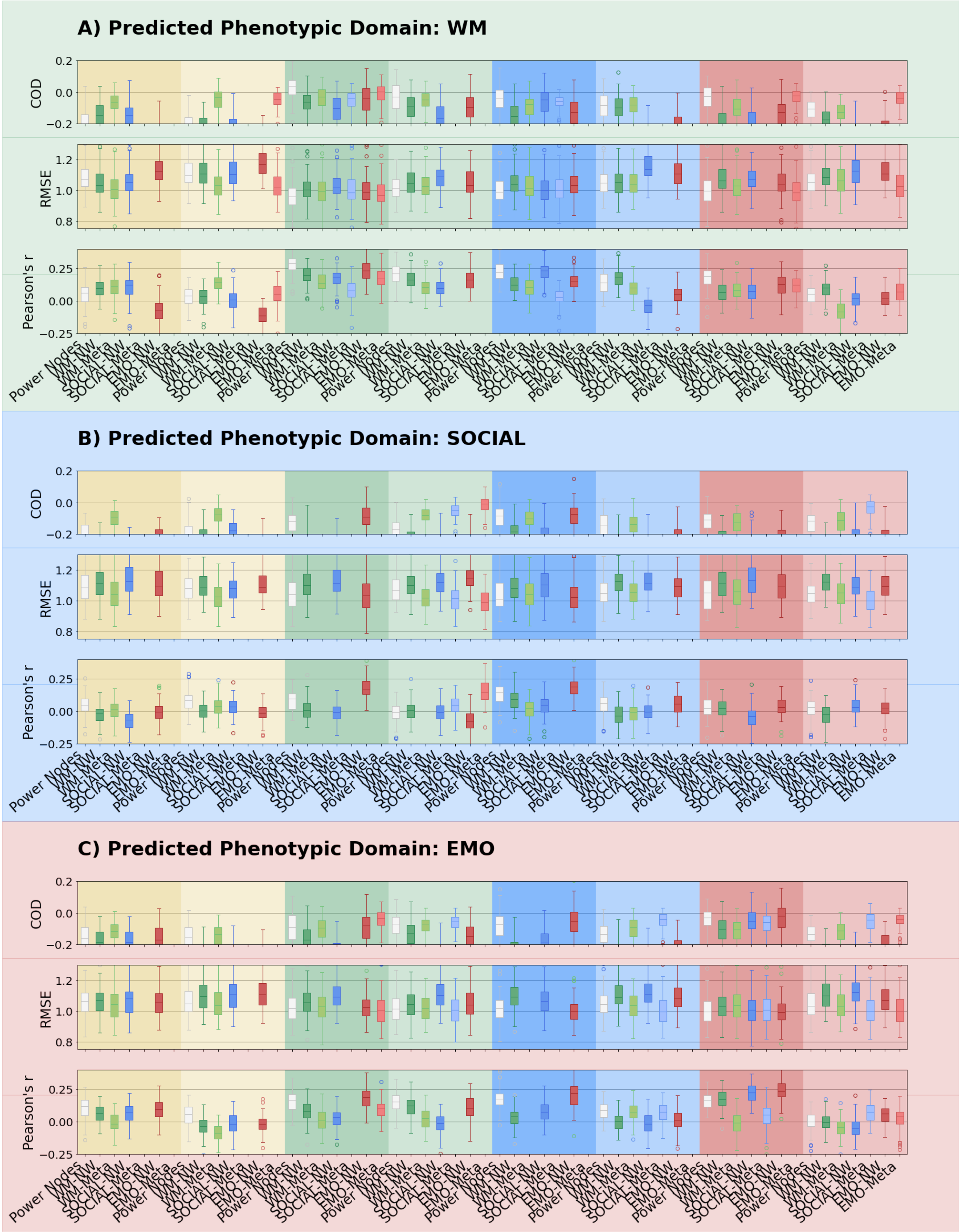


*Figure S21) CBPM – with PLS - 100 x leave-30%-out CV*

Boxplots of the distribution of prediction accuracies from CBPM – with PLS - 100 x leave-30%-out CV for WM, SOCIAL, and EMO domain, for coefficient of determination (COD) / model fit, RMSE and Pearson's r.

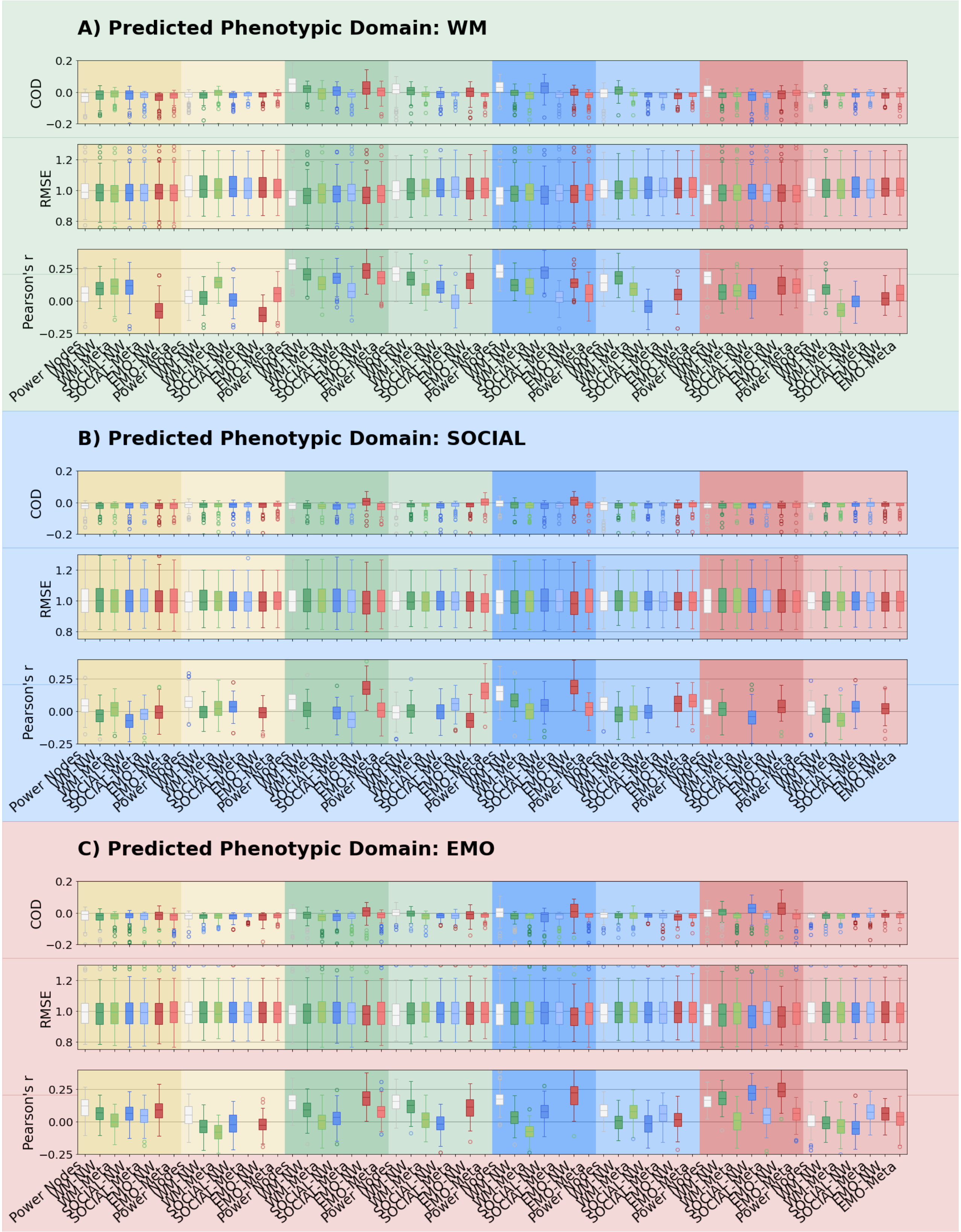


*Figure S22) CBPM – with ridge regression - 100 x leave-30%-out CV*

Boxplots of the distribution of prediction accuracies from CBPM – with ridge regression - 100 x leave-30%-out CV for WM, SOCIAL, and EMO domain, for coefficient of determination (COD) / model fit, RMSE and Pearson's r.
